# Early NK-cell and T-cell dysfunction marks progression to severe dengue in patients with obesity and healthy weight

**DOI:** 10.1101/2024.09.06.611687

**Authors:** Michaela Gregorova, Marianna Santopaolo, Lucy C. Garner, Divya Diamond, Narayan Ramamurthy, Tran Thuy Vi, Nguyet Minh Nguyen, Nguyen Lam Vuong, Eben Jones, Mike Nsubuga, Curtis Luscombe, Hoa Vo Thi My, Ho Quang Chanh, Nguyen Thi Xuan Chau, Dong Thi Hoai Tam, Duyen Huynh Thi Le, Cao Thi Tam, Paul Klenerman, Sophie Yacoub, Laura Rivino

## Abstract

Dengue is a mosquito-borne virus infection affecting half of the world’s population for which therapies are lacking. The role of T and NK-cells in protection/immunopathogenesis remains unclear for dengue. We performed a longitudinal phenotypic, functional and transcriptional analyses of T and NK-cells in 124 dengue patients using flow cytometry and single-cell RNA-sequencing. We show that T/NK-cell signatures early in infection discriminate patients who will progress to severe dengue (SD) from those who do not. In patients with overweight/obesity these signatures are exacerbated compared to healthy weight patients, supporting their increased susceptibility to SD. In SD, CD4^+^/CD8^+^ T-cells and NK-cells display increased co-inhibitory receptor expression and decreased cytotoxic capacity compared to non-SD. Furthermore, type-I Interferon signalling is downregulated in SD, suggesting defective virus-sensing mechanisms may underlie NK/T-cell dysfunction. We propose that dysfunctional “professional killer” T/NK-cells underpin dengue pathogenesis. Our findings pave the way for the evaluation of immunomodulatory therapies for dengue.

## Introduction

Dengue is caused by dengue virus (DENV), a mosquito-borne orthoflavivirus that infects an estimated 390 million people causing 300,000 severe dengue (SD) cases and 20,000 deaths yearly in tropical and subtropical countries^1^. Climate change, urbanization and human mobility are driving a rapid increase in dengue cases^2^.

DENV co-circulates as four serotypes (DENV1-4); infection with any DENV serotype can be asymptomatic or cause symptoms ranging from uncomplicated febrile illness to life-threatening SD characterized by increased vascular permeability leading to plasma leakage, potentially hypovolemic shock, organ impairment and haemorrhagic manifestations. The World Health Organization (WHO) classifies dengue cases as non-severe dengue (non-SD), non-SD with warning signs, or severe dengue (SD), the latter two requiring close clinical monitoring^3^. There are no approved therapeutics for dengue and the two licensed vaccines provide partial protective efficacy^4^. Incomplete understanding of the mechanisms underlying immune protection and progression to SD challenges the development of host-directed therapies and fully protective vaccines. Secondary infection with a different serotype is the most characterised risk factor for SD, with host immunity believed to play a central but poorly understood role in dengue pathogenesis^5^. More recently, obesity has emerged as an important risk factor^6^. The mechanisms underlying this increased risk remain unclear, but dysfunctional immune responses could be a contributing factor^7^. Alterations of the T-cell response towards pathogens as well as impaired cytotoxic functions of NK-cells during infectious disease and vaccination have been described to occur in adults and children with obesity^8–11^.

The contributions of T and NK-cells to protection/immunopathology are poorly defined. Neutralizing antibodies and CD4^+^/CD8^+^ T-cells are protective towards DENV infection^12–14^, however pre-existing cross-reactive immunity may contribute to immunopathology^12,15^. For T-cells this phenomenon, known as “original antigenic sin” (OAS), postulates that during secondary infection pre-existing cross-reactive memory T-cells with low affinity for DENV epitopes of the secondary infecting serotype may undergo suboptimal T-cell receptor triggering. This can lead to poor induction of T-cell cytotoxic functions and skewing of cytokine production towards pro-inflammatory cytokines associated with SD^15–18^. However, a study in school children shows that pre-existing TNF-a, IFN-g, and IL-2-producing DENV-specific T-cells are protective towards development of a subsequent symptomatic secondary infection, suggesting that the impact of cross-reactive T-cells in dengue is complex and the extent of occurrence of OAS in SD remains poorly understood^19^.

NK and T-cells mediate clearance of DENV-infected cells through production of anti-viral cytokines such as IFN-γ and secretion of cytotoxic granules containing perforin and granzymes. Studies by us and others, show decreased NK-cell expression of CD69, NKp30, granzyme B (GzmB), and perforin in SD compared to non-SD^20,21^. A potential impairment of immune cells mediating viral clearance during SD is consistent with studies showing the association of SD with high/prolonged viraemia and skewing of the NK-cell response from cytotoxic to cytokine-producing^22–24^. It is also consistent with genetic studies showing the association of single nucleotide polymorphisms (SNPs) in genes involved in CD8^+^ T/NK-cell cytotoxicity, namely MHC Class I Polypeptide-Related Sequence B (MICB)^25,26^, NKG2D and perforin^19^, with increased susceptibility to SD.

Here we perform an in-depth phenotypic, functional and transcriptional analyses of NK and T-cell profiles associated with disease outcomes in 124 Vietnamese dengue patients with overweight/obesity (OW/OB, N=62) or sex, age and illness phase matched healthy (HW, N=62), including 30 SD patients (**Supplementary Table S1**), at two timepoints of disease. We demonstrate an association of phenotypic and functional impairment of CD4^+^ T, CD8^+^ T and NK-cells with SD in patients enrolled prior to/at the onset of SD, with some features of immune dysfunction being exacerbated in OW/OB compared to HW patients. We propose that defective type-I IFN signalling, present across multiple cell types in SD patients, may underlie suboptimal NK and T-cell responses in SD patients. Our study provides new insights into the mechanisms underlying the progression to SD in OW/OB and HW dengue patients and paves the way for the evaluation of novel therapeutic avenues for dengue.

## Results

### Distinct T and NK-cell profiles in SD

The kinetics and phenotypic/functional features of T and NK-cell responses were investigated in peripheral blood mononuclear cells (PBMCs) from patients with SD and non-SD, with each group including patients with HW and OW/OB, at admission (T1, ≤ 5 days from fever onset) and approx. 3 days later (T2, days 6-9; **Fig.1A**). For these analyses we included a total of 146 PBMC samples from 104 dengue patients (42 patients with PBMCs at T1 + T2 and 62 patients with PBMCs available for T2 only). PBMCs were stained with antibodies targeting markers of CD4^+^/CD8^+^ T and NK-cell activation/exhaustion (CD38, HLA-DR, PD-1), proliferation (Ki-67), cytotoxicity (CD56, GPR56, GzmB, and perforin), and peripheral tissue homing [cutaneous lymphocyte-associated antigen (CLA), CCR5] and analysed by flow cytometry. In non-SD patients the frequencies of total NK-cells and NK CD56^dim^ cells, the most abundant NK-cell subset in circulation, are higher at T1 compared to T2, in line with the known early activation of NK-cells during acute viral infection; a similar trend is observed in SD HW patients but not in SD OW/OB patients (**Fig.1B**). These data suggest increasingly altered NK-cell expansion in the SD HW and OW/OB groups. In line with the known kinetics of T-cell activation, the frequencies of CD8^+^ T-cells increase from T1 to T2 consistently across all groups.

**Fig.1.**
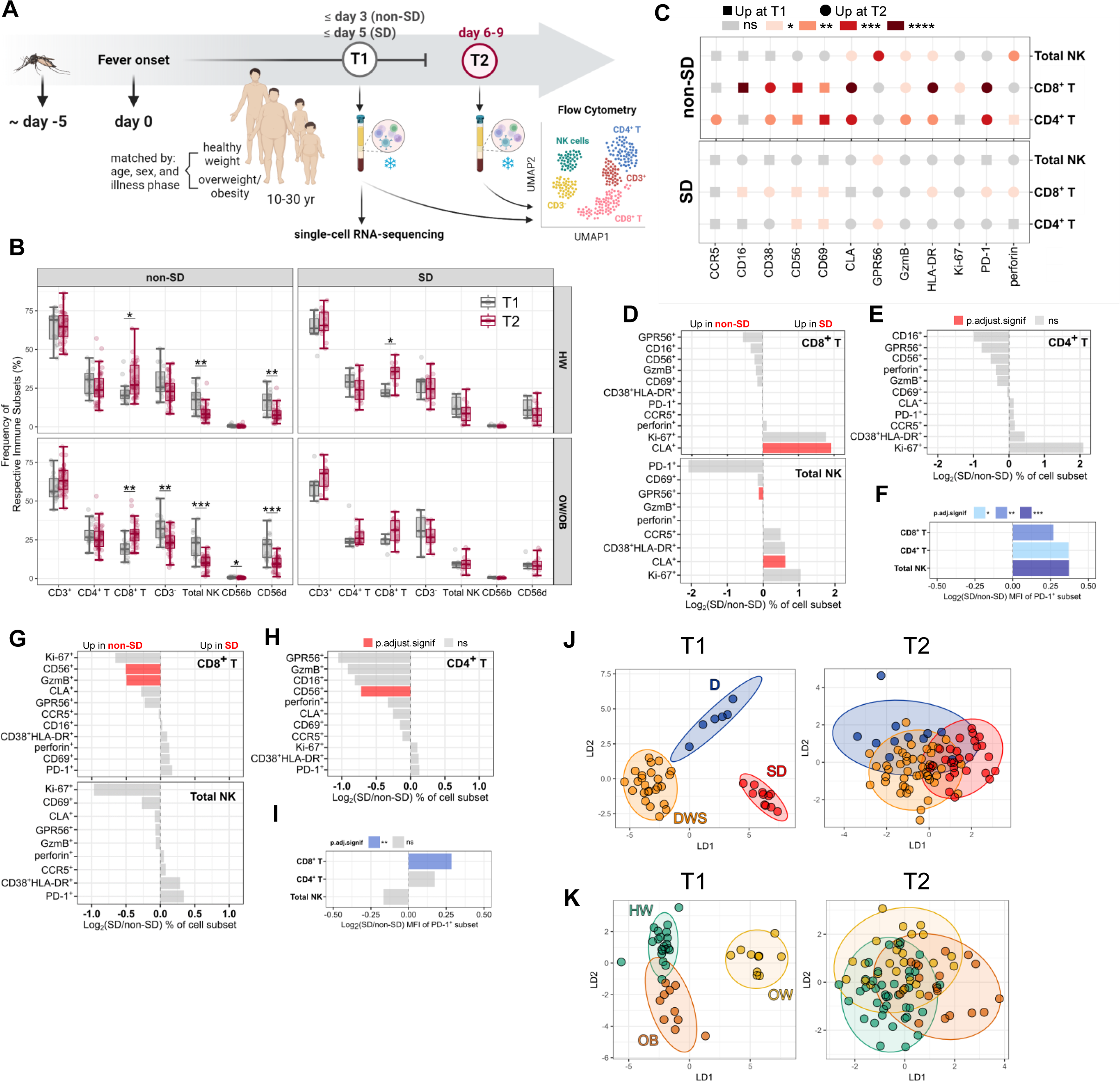
Distinct T and NK-cell profiles in SD. **(A)** Study design **(B)** Frequency of immune subsets at T1 [N=42: non-SD (N=31); SD (N=11); HW (N=21); OW/OB (N=21)] and T2 [N=104: non-SD (N=74); SD (N=27); HW (N=49); OW/OB (N=52)]. The middle line in each box represents the median with IQR. **(C)** Log_2_ ratio of MFI of selected markers in total NK, CD8^+^ and CD4^+^ T-cell subsets between T1 (N=41) and T2 (N=41) in non-SD (N=30) and SD (N=11) patients. **(D-I)** Log_2_ ration of mean abundances/PD-1^+^ MFI of cell subsets between SD and non-SD patients at T1 (N=42) **(D-F)** and T2 (N=104) **(G-I)** in CD8^+^ T, CD4^+^ T, and NK-cells. Red/blue bars indicate significance (p.adjust.signif<.05; *p.adj<.05; **p.adj<.01; ***p.adj<.001) via Wilcoxon rank sum tests. **(J)** Linear discriminant analysis at T1 (N=42) and T2 (N=84). Dots represent individual patients from dengue (D), dengue with warning signs (DWS), and SD group. Ellipses represent 95% confidence intervals. LD1 and LD2 were derived using all features shown in D-I.

We next compared the levels of expression of the analysed markers, assessed as Mean Fluorescence Intensity (MFI), within each cell type in patients with matched T1 and T2 samples (N=41 patients: N=30 non-SD; N=11 SD). The expression dynamics of most markers follows the same direction (increased/decreased at T1/T2) in both severity groups, with changes being more pronounced in non-SD patients, suggesting more dynamic immune changes in this group (**Fig.1C**). Analyses of the frequencies of CD4^+^/CD8^+^ T and NK-cells expressing the phenotypic/functional markers show increased frequencies at T1 in SD patients of CD8^+^ T and NK-cells expressing the skin homing receptor CLA, and decreased frequencies of GPR56^+^ (cytotoxic) NK-cells (**Fig.1D-E**). At T2, SD patients display decreased frequencies of cytotoxic CD4^+^/CD8^+^ T-cells (GzmB^+^/CD56^+^) compared to non-SD patients (**Fig.1G-H**). From early in infection, PD-1 expression levels are higher in SD versus non-SD in CD4^+^/CD8^+^ T and NK-cells, and these remain significantly higher at T2 for CD8^+^ T-cells, suggesting prolonged antigenic stimulation of these cells in SD (**Fig.1F, I**). PD-1 is a marker of T-cell activation and prolonged PD-1 expression marks exhausted T-cells following repetitive antigenic stimulation. Linear discriminant analysis (LDA) including all the analysed set of T/NK-cell markers discriminates at T1 patients that will progress to SD/are at the onset of SD from those that do not. Non-SD patients with warning signs map in between patients without warning signs and those with SD and are distinct from both groups (**Fig.1J**). Data at T2 can also discriminate patients based on disease severity albeit less strikingly than that at T1 (**Fig.1J**). The LDA analysis of NK/T-cell profiles also discriminates between patients who have normal weight, are overweight or have obesity suggesting an impact of patient BMI on T and NK-cell responses to DENV (**Fig.1K**). However, T/NK-cell profiles are more strongly impacted by clinical outcomes than by BMI (**Supplementary Fig.S1 A-B**). Immune profiles were similar in SD patients prior to or at the onset of SD (**Supplementary Fig.S1 C-D**). These findings suggest a progressive change of T/NK-cell features, occurring early in infection, in patients with uncomplicated dengue through to patients with SD.

### Increased PD-1^+^ CD4^+^ T-cells in SD

We next examined in more detail the phenotypic/functional profiles of CD4^+^ T-cells at admission and approx. 3 days later in non-SD and SD patients with HW and OW/OB. CD4^+^ T-cell activation defined by HLA-DR/CD38 co-expression increases from T1 to T2 and does not differ between non-SD and SD patients, nor between patients with HW or OW/OB (**Fig.2A** and **2E-F**). PD-1 expression levels also increase from T1 to T2, with a trend towards higher expression levels in SD versus non-SD patients at the earlier timepoint while expression is similar in OW/OB compared to HW patients (**Fig.2B and 2E-F**).

**Fig.2.**
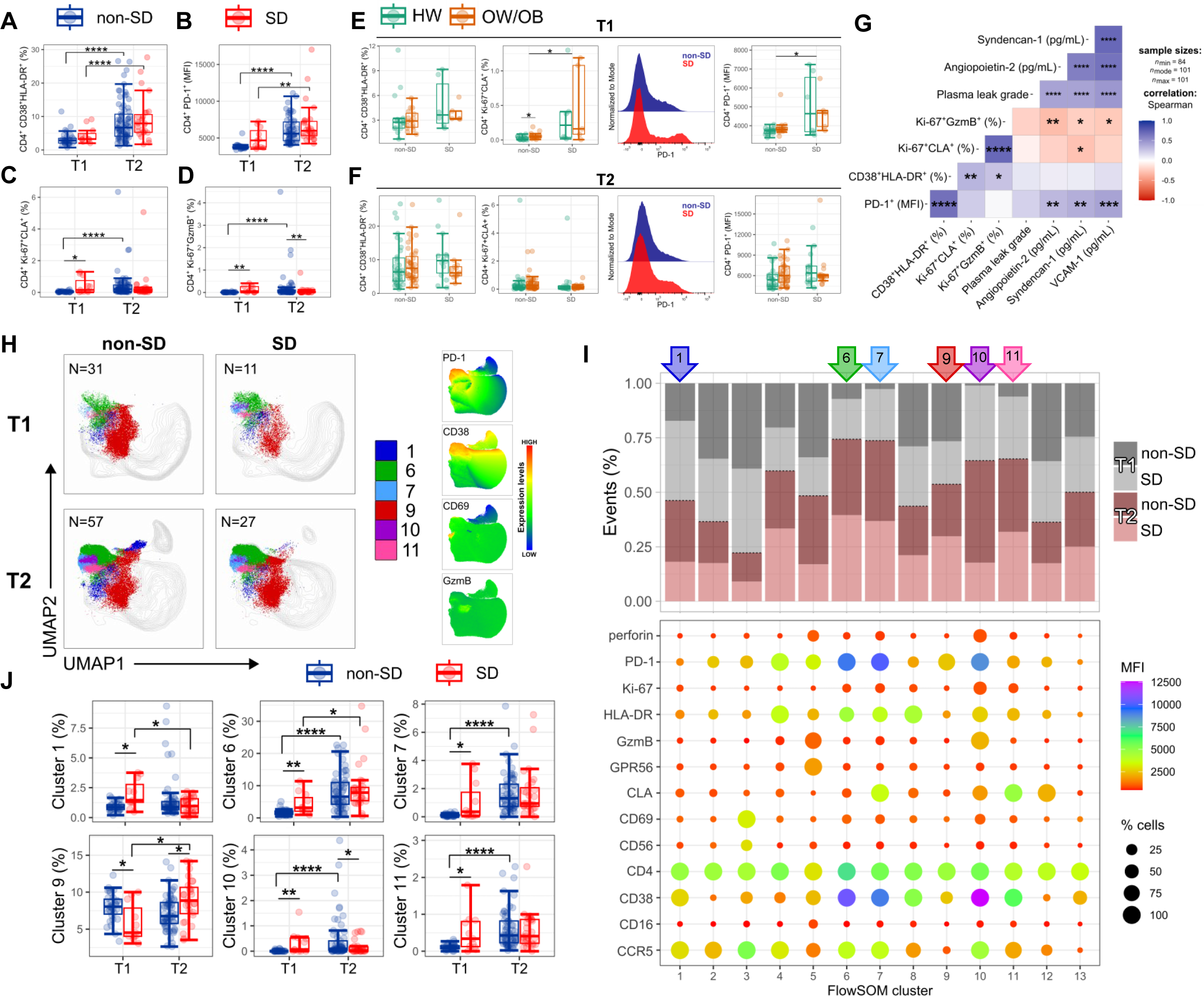
Increased PD-1^+^ CD4^+^ T-cells in SD. Percentage of CD4^+^ T-cells expressing **(A)** CD38 and HLA-DR, **(B)** Ki-67 and CLA, **(C)** Ki-67 and GzmB, and **(D)** PD-1 at T1 (N=42) and T2 (N=104) in non-SD (blue) and SD (red) patients, **(E-F)** in HW (green) and OW/OB (orange) patient groups within non-SD and SD patients at **(E)** T1 and **(F)** T2. **(G)** Correlation of CD4^+^ T-cell subsets with clinical parameters/biomarkers at T2 (N=104), using Spearman’s rank correlation test with FDR correction. **(H)** UMAP plots with FlowSOM clusters (1,6,7,9-11) visualised in non-SD and SD patient groups at T1 (N=42) and T2 (N=84) and expression levels of PD-1, CD38, CD69, and GzmB. **(I)** Stacked bar chart showing the frequency of each cluster in specific groups with bubble graph representing MFI levels (colour) and cell frequencies (size). **(J)** Frequency of FlowSOM clusters (1,6,7,9-11) at T1 (N=42) and T2 (N=84) within non-SD and SD patient groups. The middle line in each box represents the median with IQR. Error bars represent max/min value +/- 1.5*IQR. *p<.05; **p<.01; ***p<.001; ****p<.0001 by one-way ANOVA with Benjamini-Hochberg correction.

Ki-67 and CLA co-expression can be used as a proxy for DENV-specific T-cells^27,28^, hence Ki-67^+^CLA^+^ are herein also named *bona fide* DENV-specific T-cells. In non-SD patients, the frequencies of Ki-67^+^CLA^+^ CD4^+^ T-cells increased from T1 to T2, mirroring CD4^+^ T-cell activation. However, in SD patients CD4^+^ Ki-67^+^CLA^+^ T-cells are present at higher frequencies early at T1 compared to non-SD patients, and then fail to expand further, suggesting different kinetics of expansion of *bona fide* DENV-specific CD4^+^ T-cells in SD patients (**Fig.2C**). The frequencies of Ki-67^+^GzmB^+^ CD4^+^ T-cells follow similar trends (**Fig.2D**). Non-SD OW/OB patients display a modest increase in Ki-67^+^CLA^+^ DENV-specific CD4^+^ T-cells at T1 compared to HW non-SD (**Fig.2E-F**), similarly to what is observed in SD patients. Otherwise, expression of the analysed markers appears similar across BMI groups.

Plasma levels of the markers of endothelial dysfunction angiopoietin-2, syndecan-1, and VCAM-1 show a positive moderate correlation with PD-1 expression in CD4^+^ T-cells (**Fig.2G**, shown for T2). Conversely, these endothelial dysfunction-related markers correlated negatively with the frequency of Ki-67^+^GzmB^+^ and Ki-67^+^CLA^+^ CD4^+^ T-cells. These data suggest a potential protective role of responding cytotoxic and *bona fide* DENV-specific CD4^+^ T-cells and a detrimental role of PD-1^+^ CD4^+^ T-cells in dengue.

To better understand the combinatorial expression of phenotypic/functional markers in CD4^+^ T-cells, we performed unsupervised dimensionality reduction analysis using uniform manifold approximation and projection (UMAP) and the FlowSOM clustering (**Fig.2H-J**). FlowSOM analyses of concatenated flow cytometry standard (FCS) files at T1/T2 from a total of 126 patient samples identifies 13 phenotypically distinct CD4^+^ T-cell clusters based on the expression of the analysed markers. The frequencies of each cluster within non-SD and SD patients at T1/T2, and expression of markers in each cluster are shown in **Fig.2I**. The six clusters that were most distinctive in non-SD and SD patients at T1 and T2 are characterised by high expression of PD-1, CD38, CD69 and GzmB (**Fig.2H** right panel). Clusters 1, 6, 7 and 10-11 are highly expressed in SD versus non-SD patients at T1 and include activated CD4^+^ T-cells with low cytotoxic potential (cluster 1, 6, 7) as well as high PD-1 and CCR5 expression (clusters 6, 7). Moreover, CD4^+^ T-cells in cluster 7 display increased expression of CLA and Ki-67, suggesting this cluster of cells could represent *bona fide* DENV-specific CD4^+^ T-cells. In summary, manual gating and unsupervised analyses show increased CD4^+^ T-cell expression of PD-1 in SD compared to non-SD patients, and expansion of PD-1^+^ CD4^+^ T-cell populations with low cytotoxic potential with these CD4^+^ T-cell features correlating with endothelial markers of SD.

### Altered PD-1 and GzmB expression in CD8^+^ T-cells in SD

Similarly to CD4^+^ T-cells, CD8^+^ T-cell activation increases from T1 to T2 but does not differ in patients across disease severities (**Fig.3A**). In non-SD the frequencies of cytotoxic GzmB^+^ CD8^+^ T-cells increase from T1 to T2, mirroring CD8^+^ T-cell activation. However, in SD patients the frequencies of GzmB^+^ CD8^+^ T-cells appear to uncouple from CD8^+^ T-cell activation and are decreased compared to non-SD patients at T2 (**Fig. 3B).** Ki-67^+^CLA^+^ (*bona fide* DENV-specific) CD8^+^ T-cells display similar expansion kinetics to their CD4^+^ T-cell counterparts, with SD patients displaying a higher frequency of these cells at T1 compared to non-SD patients, and an opposite trend is observed at the later time point (**Fig. 3C**). PD-1 levels also increase in time from T1 to T2 with SD patients displaying significantly higher expression levels compared to non-SD patients at both time points (**Fig. 3D**). Moreover, at T2, we detected decreased GzmB expression in total CD8^+^ T-cells as well as in PD-1^+^ CD8^+^ T-cells in SD patients, suggesting an impairment in their cytotoxic function which could potentially contribute to inefficient virus clearance known to occur in SD (**Fig. 3B and 3E**). Data stratified by patient BMI status and disease severity shows similar frequencies of activated and GzmB^+^ CD8^+^ T-cells in HW and OW/OB patients, but a trend towards increasingly higher PD-1 expression levels from non-SD HW, non-SD OW/OB, SD HW and SD OW/OB patients at both timepoints, suggesting that in dengue PD-1 expression in CD8^+^ T-cells may be exacerbated by high BMI (**Fig 3F-G**). PD-1 expression in CD8^+^ T-cells shows a strong positive correlation with plasma leakage grade, markers of endothelial dysfunction and with the dengue severity-related markers (ferritin and CXCL-10/IP-10) at both time points (**Fig.3H** shown for T2; **Fig 3I-J**: angiopoietin-2 and VCAM-1, shown for T1 and T2). Conversely, GzmB expression in CD8^+^ T-cells correlates negatively with plasma leakage and endothelial dysfunction, suggesting a protective role of cytotoxic CD8^+^ T-cells in dengue. Moreover, the frequency of activated HLA-DR^+^CD38^+^ CD8^+^ T-cells inversely correlate with the plasma levels of leptin, suggesting that obesity could have an impact on CD8^+^ T-cell activation during dengue infection.

**Fig.3.**
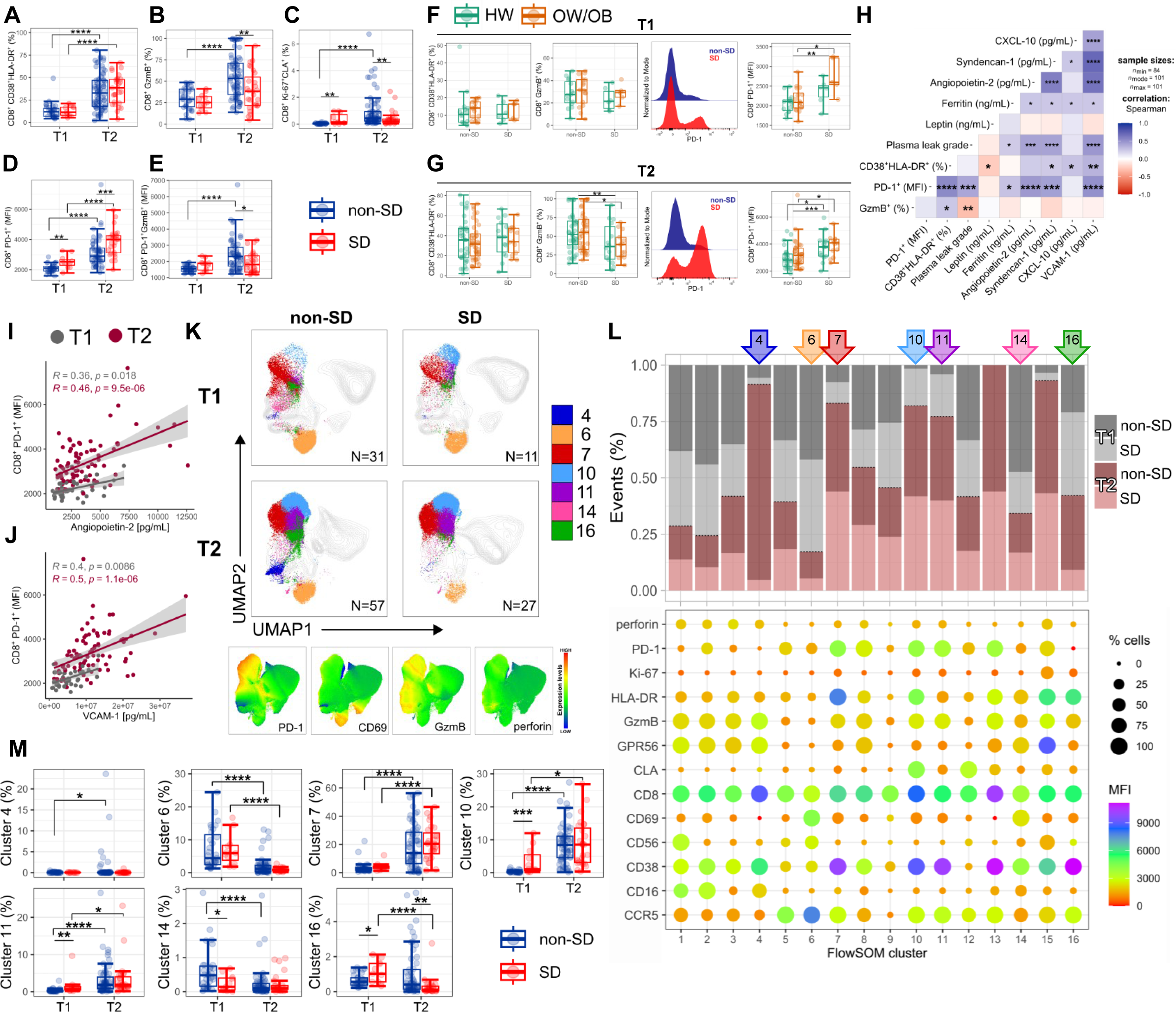
Altered PD-1 and GzmB expression in CD8^+^ T-cells in SD patients. Percentage of CD8^+^ T-cells expressing **(A)** CD38 and HLA-DR, **(B)** GzmB, **(C)** Ki-67 and CLA, **(D)** PD-1 and **(E)** GzmB on PD-1^+^ CD8^+^ T-cells at T1 (N=42) and T2 (N=104) in non-SD and SD patients and in HW (green) and OW/OB (orange) patient groups within non-SD and SD patients at **(F)** T1 and **(G)** T2. **(H)** Correlation of CD8^+^ T-cell subsets with clinical parameters/biomarkers at T2 (N=104), using Spearman’s rank correlation test with BH correction. Single correlations of CD8^+^ T-cells expressing PD-1 with plasma levels of **(I)** Angiopoietin-2 and **(J)** VCAM-1 at T1 (N=42) and T2 (N=104). **(K)** UMAP plots with FlowSOM clusters (4,6,7,10,11,14,16) visualised in non-SD and SD patient groups at T1 (N=42) and T2 (N=84) and expression levels of PD-1, CD69, GzmB, and perforin. **(L)** Stacked bar chart showing the frequency of each cluster in specific groups and bubble graph represents MFI levels (colour) and cell frequencies (size). **(M)** Frequency of FlowSOM clusters (4,6,7,10,11,14,16) at T1 (N=42) and T2 (N=84) within non-SD and SD patient groups. The middle line in each box represents the median with IQR. Error bars represent max/min value +/- 1.5*IQR. *p<.05; **p<.01; ***p<.001; ****p<.0001 by one-way ANOVA with Benjamini-Hochberg correction.

Unsupervised analyses using UMAP and FlowSOM clustering of concatenated FCS files at T1/T2 from 126 patient samples, reveals 16 distinct CD8^+^ T-cell clusters based on the combinatorial expression of the analysed markers (**Fig.3K-M**). The clusters with the most distinct representation between non-SD and SD patient groups at T1 and T2 fall within areas of highest expression of PD-1, CD69, GzmB and perforin (**Fig.3K** bottom panel). Clusters 10, 11 and 16 are present at higher frequencies in SD compared to non-SD patients at T1 and contain CD8^+^ T-cells with high PD-1 and CCR5 expression. Cells from cluster 10 are CLA^+^Ki-67^+^ and may represent *bona fide* DENV-specific CD8^+^ T-cells. Cluster 4 is detected only in non-SD patients, although in a minor proportion of these patients (N=7/42), and contains moderately activated cells, with high cytotoxic potential (GzmB^+^, GPR56^+^) and low PD-1 and CCR5 expression. Clusters 6 and 7 contain cells that are respectively increased at T1 and T2, suggesting they represent T-cell populations at early and late differentiation stages. Accordingly, CD8^+^ T-cells in cluster 6 cells express high levels of the early activation marker CD69 and CCR5 while those in cluster 7 express the activation markers HLA-DR/CD38, PD-1 and display increased cytotoxic potential. Collectively these data demonstrate decreased PD-1 and GzmB expression in CD8^+^ T-cells from SD patients, which strongly correlates with clinical markers of dengue disease severity.

### Altered DENV NS3-specific T-cell response in SD

To address whether disease severity and BMI associate with altered DENV-specific T-cell responses we evaluated CD4^+^ and CD8^+^ T-cell responses to overlapping peptides spanning the immunodominant NS3 protein^29^. Intracellular cytokine staining (ICS) was performed to measure production of IFN-γ, TNF-α, IL-2, MIP-1β, and CD107a, an indirect marker of degranulation by flow cytometry (**Fig.4A**). To investigate the phenotypic features of DENV-specific T-cells we co-stained cells with antibodies targeting markers of T-cell differentiation (CD95) and activation/exhaustion (PD-1). For these analyses we selected patients with secondary DENV-2 infection. The frequencies of DENV2 NS3-specific CD4^+^ and CD8^+^ T-cells, defined as cytokine^+^ and/or CD107a^+^ T-cells, are higher in SD compared to non-SD patients at T1, with an opposite trend at the later time point, similarly to the *bona fide* DENV-specific T-cells (**Fig.4B**). To investigate whether there was a skewed T-cell response to DENV serotypes potentially encountered during the primary infection (OAS), driven by expansion of pre-existing memory T-cells, we tested recognition of NS3 peptide pools from all 4 DENV serotypes. Cytokine production by CD4^+^ and CD8^+^ T-cells upon recognition of NS3 peptide pools from DENV1-4 serotypes is similar, but responses are overall lower in SD compared to non-SD patients at T2 (**Fig.4C and supplementary Fig.S2**). These data suggest broad cross-reactive T-cell recognition of NS3 peptides from all 4 DENV serotypes with no preferential skewing for a specific serotype, and increased magnitude of DENV-specific T-cell responses in non-SD patients at the time of viral clearance.

**Fig.4.**
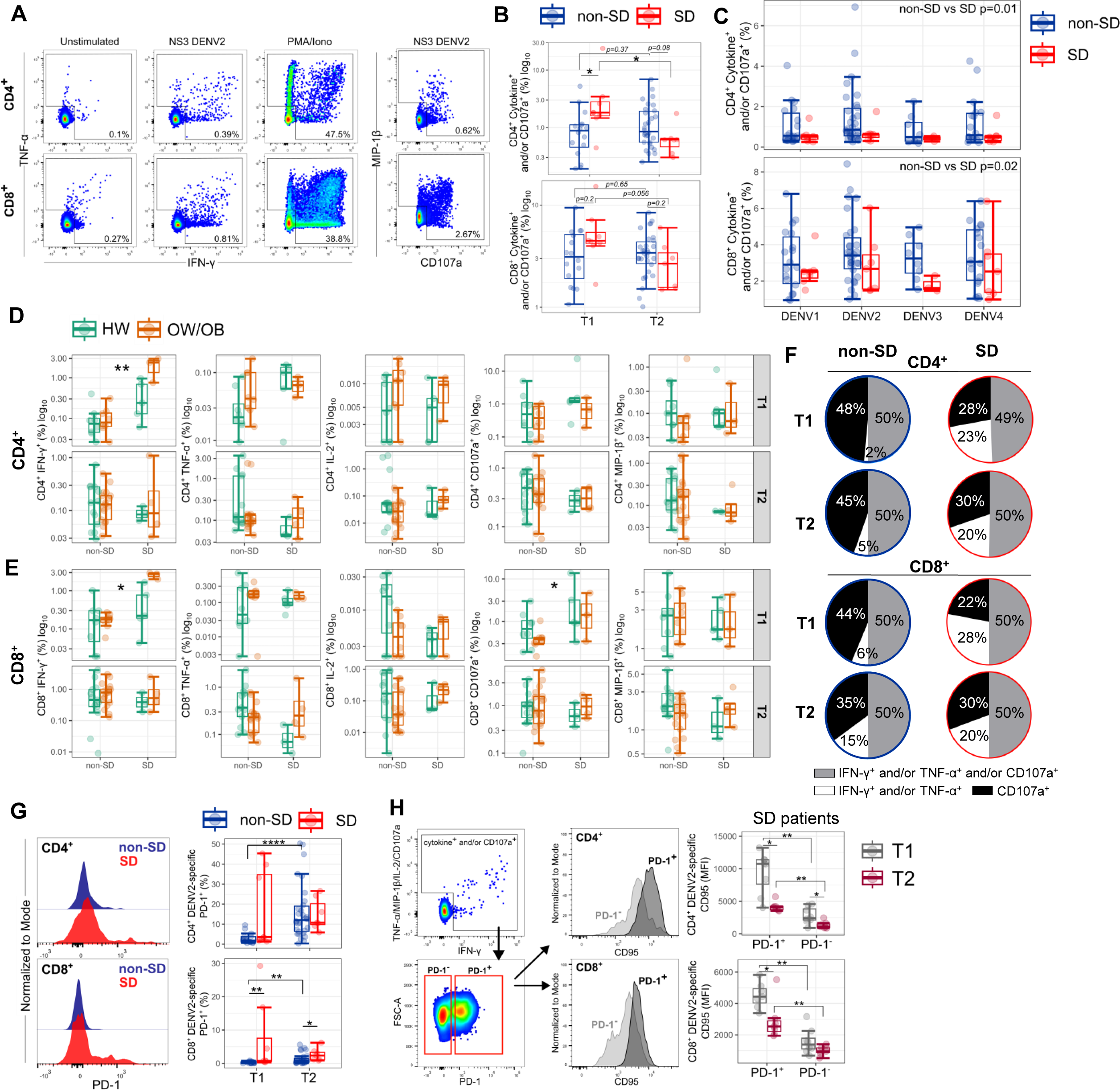
Altered DENV NS3-specific T-cells in SD. **(A)** Representative flow cytometry plots showing IFN-γ, TNF-α, MIP-1β, and CD107a production by CD4^+^ and CD8^+^ T-cells from non-SD patient (day 6 of fever) after stimulation with NS3 DENV2 peptides, PMA/ionomycin, and unstimulated condition (DMSO). **(B)** CD4^+^ and CD8^+^ T-cell responses shown as log10 of cytokine^+^ (IFN-γ and/or TNF-α and/or IL-2 and/or MIP-1β) and/or CD107a ^+^ cells at T1 (N=24) and T2 (N=37) after stimulation with NS3 DENV2 peptides. **(C)** CD4^+^ and CD8^+^ T-cell responses shown as cytokine^+^ and/or CD107a^+^ at T2 (N=37) after stimulation with NS3 DENV1-4 peptide pools. **(D,E)** Single cytokine response by **(D)** CD4^+^ and **(E)** CD8^+^ T-cells in HW or OW/OB non-SD and SD patients at T1 (N=24) and T2 (N=37) following NS3 DENV2 peptide stimulation. **(F)** Cytokine^+^ and/or CD107a^+^ T-cells were divided into three groups: degranulation only (CD107a), degranulation and cytokine production (IFN-γ and/or TNF-α and/or CD107a), and cytokine production only (IFN-γ and/or TNF-α). Pie charts show the percentages of the cytokine^+^ and/or CD107a^+^ cells to total responding T-cells from non-SD and SD patients. The average percentages are sum from all individuals and divided by number of cases. **(G)** Representative histogram and boxplots of PD-1 expression by DENV2-specific CD4^+^ and CD8^+^ T-cells at T1 (N=24) and T2 (N=37). **(H)** Gating strategy and representative flow cytometry plots of CD95 and PD-1 expressing T-cells and comparison between CD95 MFI levels in PD-1^+^ and PD-1^-^ CD4^+^ and CD8^+^ T-cells in SD patients. The middle line in each box represents the median with IQR. Error bars represent max/min value +/- 1.5*IQR. *p<.05; **p<.01; ***p<.001; ****p<.0001 by one-way ANOVA with Benjamini-Hochberg correction.

We next investigated whether T-cell cytokine profiles differ across patient groups (**Fig.4D-E**). DENV2-specific T-cells are mainly monofunctional (i.e. expressing a single cytokine/function) with a minor proportion of polyfunctional cells expressing 2-5 functions (**Supplementary Fig.S3**). We therefore focused on analysis of T-cells producing single functions. Early in infection, at T1, DENV2-specific CD4^+^ T-cells mainly produce IFN-γ and CD107a with a stepwise increase of IFN-γ production observed from non-SD to SD HW and SD OW/OB patients (**Fig 4D**, top panel). DENV2-specific CD8^+^ T-cells display a similar stepwise increase of IFN-γ production from non-SD to SD HW and SD OW/OB patients at T1 (**Fig.4E**, top panel). At this timepoint, SD patients also display increased frequencies of degranulating CD107a^+^ NS3-specific CD8^+^ T-cells. At T2, IFN-γ/CD107a production by DENV2-specific CD4^+^ and CD8^+^ T-cells was comparable across groups (**Fig.4D-E**, bottom panels). Production of IL-2, TNF-α and MIP-1β by NS3-specific CD4^+^ and CD8^+^ T-cells was also largely similar across groups at both timepoints. These findings are in line with the higher frequencies of *bona fide* DENV-specific T-cells (**Fig.2C/3C**) and total cytokine^+^/CD107a^+^ T-cells (**Fig.4B**) at T1 in SD compared to non-SD patients.

In line with previous studies^16^, at T1 DENV(NS3)-specific T-cells in SD patients contained higher percentages of cytokine-producing/CD107a-negative and lower percentages of cytokine-negative/CD107a-positive cells, suggesting a skewing of T-cell responses towards cytokine production in SD and conversely towards cytotoxicity in non-SD patients (**Fig.3F**). As SD patients display higher viraemia early in infection^20,30^, we asked whether T-cells may be undergoing excessive antigen-driven activation leading to antigen-induced cell death. A higher proportion of NS3 DENV2-specific CD4^+^ and CD8^+^ T-cells expressed PD-1 at T1 in SD patients, while these cells are largely absent in non-SD patients (**Fig.4G**). PD-1^+^ NS3 DENV2-specific CD4^+^ and CD8^+^ T-cells from SD patients expressed high levels of the death receptor CD95 compared to their PD-1^-^ counterparts, suggesting these cells are undergoing cell death, possibly due to excessive antigenic stimulation. At T2, CD95 levels of PD-1^+^ cells are significantly decreased from T1, but they remain higher compared to PD-1^-^ cells (**Fig.4H**). Collectively, these data suggest that SD patients display higher frequencies of NS3 DENV-specific T-cells early in infection, which are skewed towards cytokine production, express PD-1 and CD95 and may be prone to apoptosis.

### Elevated T-cell co-inhibitory receptors in SD

We next asked whether T-cells in SD patients express other co-inhibitory receptors beyond PD-1. To this end we analysed expression of 5 co-inhibitory receptors associated with T-cell exhaustion (CTLA-4, LAG-3, TIM-3, PD-1, and TIGIT)^31^. At T1, SD patients display significantly increased frequencies of TIGIT^+^ and TIM-3^+^ CD4^+^ T-cells and LAG-3^+^ CD8^+^ T-cells and a trend towards increased frequencies of CD4^+^/CD8^+^ T-cells expressing all co-inhibitory receptors analysed (**Fig.5A-B**). Similar to what we observed for PD-1, the frequencies of CD4^+^/CD8^+^ T-cells expressing these inhibitory receptors correlates positively with endothelial dysfunction and severity-related plasma markers (syndecan-1, VCAM-1, and ferritin) (**Fig.5C-D**). In SD patients there was a larger frequency of CD4^+^ and CD8^+^ T-cells co-expressing multiple co-inhibitory receptors compared to non-SD patients, with TIGIT and PD-1 being the most highly expressed (**Fig.5E**), suggesting these T-cells may be exhausted. To gain further insights into the features of T-cells expressing multiple co-inhibitory receptors we established a flow cytometry panel which included 6 co-inhibitory receptors (CTLA-4, LAG-3, TIM-3, PD-1, TIGIT and LILRB1), activation/proliferation markers (ICOS, CD25, Ki-67) and markers for T-cells (CD3, CD4, CD8), NK-cells (CD16, CD56) and Tregs (CD25, FOXP3). UMAP and FlowSOM clustering analyses identified 15 distinct CD4^+^ and CD8^+^ T-cell clusters; clusters showing significant differences in SD versus non-SD patients are colour-coded in the UMAP (**Fig.5F**: CD4^+^ T clusters 13-15; **Fig.5G**: CD8^+^ T clusters 12-14). CD4^+^ T-cells in clusters 13-15 are largely detectable only in SD patients. Cluster 13 contains proliferating ICOS^+^ CD4^+^ T-cells co-expressing all 6 co-inhibitory receptors as well as CD16 and CD56. Cells in cluster 14 co-express 4 inhibitory receptors although at lower levels compared to cluster 13 and they lack expression of Ki-67. Cluster 15 contains proliferating CD4^+^ CD25^+^FOXP3^+^ regulatory T-cells (Tregs) expressing TIGIT and CD56 (**Fig.5F**). Similarly, SD patients display increased levels of non-proliferating CD8^+^ T-cells expressing co-inhibitory receptors TIGIT, TIM-3, and PD-1 or ICOS, LILRB1, CTLA-4, LAG-3, PD-1 and TIGIT (**Fig.5G**, respectively clusters 13 and 14). Interestingly CD8^+^ T-cells in the latter cluster also express FOXP3, suggesting they may represent CD8^+^ Tregs.

**Fig.5.**
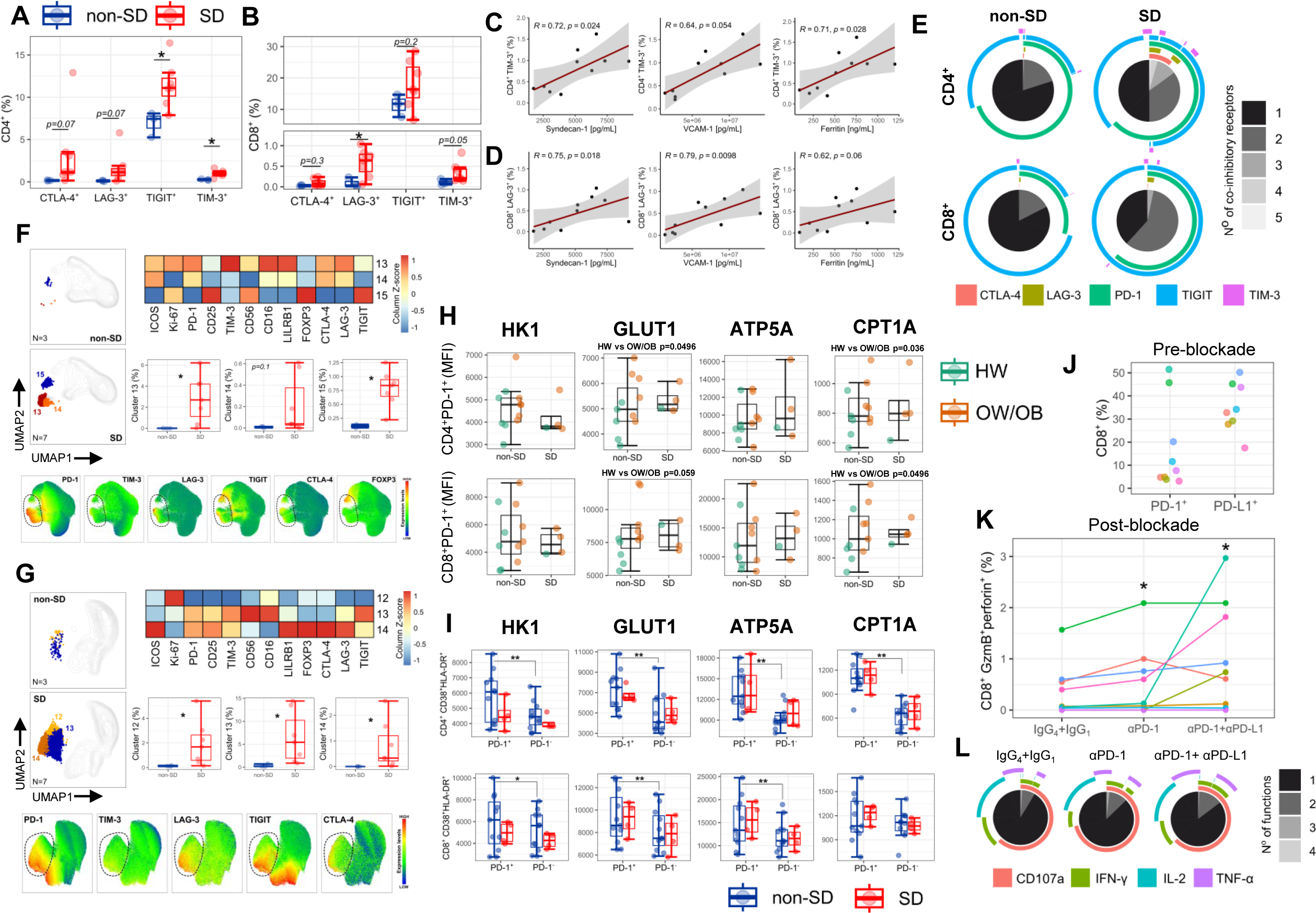
Elevated T-cell co-inhibitory receptors in SD. Expression levels of CTLA-4, LAG-3, TIGIT, and TIM-3 by **(A)** CD4^+^ and **(B)** CD8^+^ T-cells at T1 (N=10) in non-SD and SD patients. Correlation of **(C)** CD4^+^ TIM-3^+^ and **(D)** CD8^+^ LAG-3^+^ T-cells with plasma biomarkers (Syndecan-1, VCAM-1, and ferritin). **(E)** Pie charts showing the number of co-inhibitory receptors simultaneously exhibited by CD4^+^ and CD8^+^ T-cells in non-SD and SD patients at T1 (N=11). The different shades of grey represent the range of 1 to 5 co-inhibitory receptors and outer arcs indicate the specific co-inhibitory receptors detected defined by Boolean gating. **(F,G)** UMAP plots with FlowSOM clusters of **(F)** CD4^+^ and **(G)** CD8^+^ T-cells at T1 (N=11) in non-SD and SD patients. Expression levels of markers (heatmap and UMAP) and frequency of clusters in non-SD and SD group. **(H)** Expression levels of metabolic markers (HK1, GLUT1, ATP5A, and CPT1A) in CD4^+^ and CD8^+^ PD-1^+^ T-cells in HW (green) and OW/OB (orange) patient groups within non-SD and SD patients with day 5-8 of fever and **(I)** the difference between PD-1^+^ and PD-1^-^ subsets (N=17). **(J)** Expression levels of PD-1 and PD-L1 by CD8^+^ T-cells in non-SD patients (N=9). **(K)** Frequency of GzmB^+^perforin^+^ CD8^+^ T-cells after stimulation with NS3 DENV peptides and with/without blocking antibodies. **(L)** Pie charts showing the number of functions simultaneously exhibited by CD8^+^ T-cells following NS3 DENV peptide stimulation with/without blocking antibodies. The different shades of grey represent the range of 1-4 functions, the outer arcs indicate the specific functions defined by Boolean gating. The middle line in each box represents the median with IQR. Error bars represent max/min value +/- 1.5*IQR. *p<.05; **p<.01; ***p<.001; ****p<.0001 by one-way ANOVA with Benjamini-Hochberg correction.

Collectively, these data show higher frequencies in SD compared to non-SD, of responding (activated and/or proliferating) CD4^+^ and CD8^+^ T-cells co-expressing inhibitory receptors, with highest PD-1 and TIGIT expression, as well as Treg populations.

T-cell function is closely linked with cellular metabolism and the latter is shown to be altered in obesity. We therefore asked whether T-cells expressing co-inhibitory receptors have altered metabolic profiles in SD and in OW/OB, using PD-1 as a representative marker. For these analyses we selected a total of 17 patients; all except one patient had a secondary DENV2 infection (non-SD: N=11; SD: N=6 including N=8 HW and N=9 overweight/obesity; illness days 5-8). The metabolic profiles of PD-1^+^/PD-1^-^ CD4^+^ and CD8^+^ T-cells were assessed by measuring expression by flow cytometry of four key metabolic enzymes/components involved in ATP biosynthesis (ATP5A), fatty acid oxidation, FAO (CPT1A), and glycolysis (HK1, and GLUT1). PD-1^+^ CD4^+^ and CD8^+^ T-cells from OW/OB patients display increased expression levels of GLUT1 and CPT1A compared to their HW counterparts (**Fig.5H**), with levels being similar in non-SD and SD patients. PD-1^+^ CD4^+^ and CD8^+^ T-cells display elevated metabolic activity compared to their PD-1^-^ counterparts, based on expression of HK1, GLUT1, ATP5A and CPT1A, with differences between statistically significant for non-SD but not for SD patients (**Fig.5I**). Similar results were obtained using the SCENITH method (Single Cell Energetic metabolism by profiling translation inhibition)^32,33^**(Supplementary Fig.S4)**. These data suggest that PD-1^+^ CD4^+^ and CD8^+^ T-cells are engaging in glycolysis, FAO and ATP biosynthesis to support their metabolic demands, more so in OW/OB compared to HW patients.

We next asked whether PD-1 expression is playing a role in inhibiting anti-viral T-cell effector functions in dengue, specifically their cytotoxic potentially which appears to be impaired. To address this, we performed an overnight stimulation with NS3 DENV peptides (matched to the serotype of infection) in the presence of anti-PD-1 and/or anti-PD-L1 blocking antibodies or an isotype control and measured cytokine production and GzmB/perforin production of CD4^+^ and CD8^+^ T-cells. Prior to anti-PD(L)1 blockade, we evaluated *ex vivo* CD8^+^ T-cells expression of PD-1 and PD-L1 (**Fig.5J**). After stimulation and blockade, we measured expression of cytotoxic mediators and cytokine production. While the effects on DENV-specific T-cells were difficult to evaluate due to the limiting number of these cells, overall PD-1 and anti-PD-1/PDL-1 blockade led to increased frequencies of GzmB^+^ perforin^+^ CD8^+^ T-cells, with variation across patients (**Fig.5K**). While PD-1 and PD-1/PDL-1 blockade did not affect the frequencies of CD8^+^ T-cells producing each analysed cytokine, there was a trend towards increased T-cell polyfunctionality defined as co-expression of CD107a, IFN-γ and TNF-α (**Fig.5L**).

### Impaired NK-cell responses in SD

NK-cell function is governed by the balance of signals from cell surface activating and inhibitory receptors^34^. We therefore assessed NK-cell expression of activating (NKG2D, NKp46) and inhibitory receptors (LILRB1, NKG2A, PD-1, PDL-1, TIGIT and LAG-3) as well as markers of activation, proliferation (CD69, Ki-67), differentiation (CD57, NKG2C) and cytotoxic potential (GzmB, perforin) in samples from 23 patients (non-SD: N=11; SD: N=12) at T1. To determine the features of NK-cells that are responding to DENV infection, we analysed proliferating Ki-67^+^ NK-cells, herein defined as responding NK-cells (**Fig.6A**). Data is shown for total responding NK-cells (findings were similar for NK CD56^dim^ and NK CD56^bright^ cells, data not shown). In line with our previous findings, responding NK-cells in dengue infection are predominantly immature CD57^-^NKG2C^-^ cells^35^, with this being more pronounced in SD patients (**Fig.6B)**. Interestingly, SD patients display lower frequencies of CD57^+^NKG2C^+^ “memory” NK-cells. Responding NK-cells from SD patients are less activated (CD69) and displayed decreased cytotoxic potential (GzmB, perforin) and decreased expression of activating receptors NKG2D and NKp46, compared to those from non-SD patients (**Fig.6C**). Conversely, the expression levels of NK-cell co-inhibitory receptors LILRB1, NKG2A, PD-L1, TIGIT and PD-1 and the frequencies of LAG-3^+^ NK-cells are higher in SD compared to non-SD patients. Collectively, these data suggest a potential impairment in the NK-cell response in SD patients.

**Fig.6.**
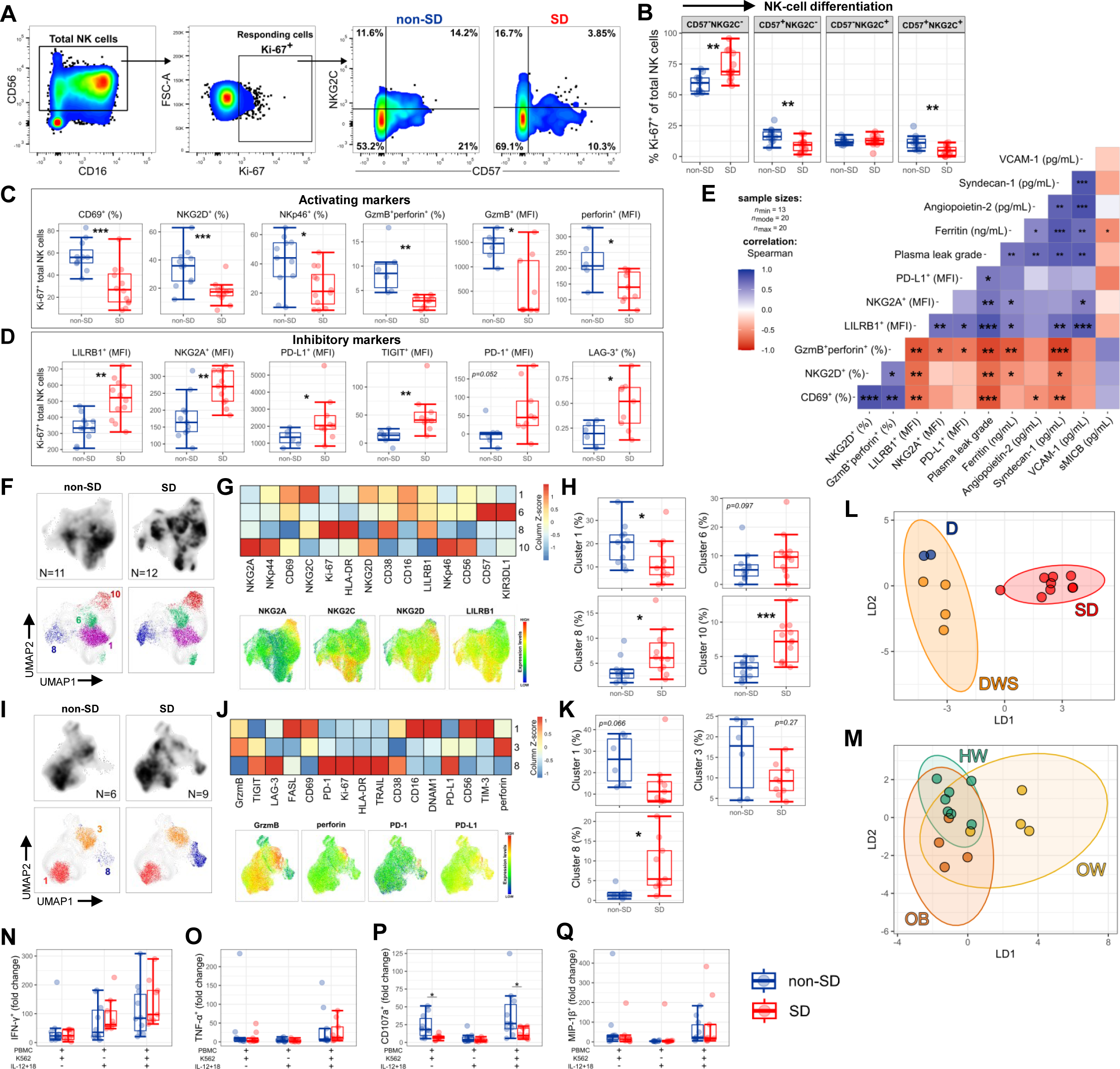
Impaired NK-cell responses in SD. **(A)** Gating strategy and representative flow cytometry plots of CD57 and NKG2C staining in non-SD and SD samples at T1. **(B)** Frequency of Ki-67^+^ total NK-cells with differential expression of CD57 and NKG2C in non-SD (N=11) and SD (N=12) patients. (**C,D**) Expression of activating and inhibitory receptors by Ki-67^+^ total NK-cells. **(E)** Correlation of NK-cell markers with clinical parameters/biomarkers using Spearman’s rank correlation test with Benjamini-Hochberg correction. **(F,I)** UMAP plots with colour coded Phenograph clusters of total NK-cells in non-SD and SD patients. **(G,J)** Expression levels of markers (heatmap and UMAP) and **(H,K)** frequency of clusters in non-SD and SD group. **(L,M)** Linear discriminant analysis at T1 (N=23). Dots represent individual patients from dengue (D), dengue with warning signs (DWS), and SD **(L)** and HW, OW, and OB group **(M)**. Ellipses represent 95% confidence intervals. LD1 and LD2 were derived using all features shown in B-D, H, K, and expression levels of KIR3DL1, NKp44, and TIM-3. **(N-Q)** Expression of cytokines (IFN-γ, TNF-α, MIP-1β) and CD107a by NK-cells after stimulation with PBMC+K562, PBMC+IL-12+IL-18, and PBMC+K562+IL-12+IL-18. Data are shown as fold change from the baseline condition (PBMC only). The middle line in each box represents the median with IQR. Error bars represent max/min value +/- 1.5*IQR., *p<.05; **p<.01; ***p<.001; ****p<.0001 by Wilcoxon test or one-way ANOVA with Benjamini-Hochberg correction.

Plasma leakage grade, endothelial dysfunction (angiopoietin-2, sydnecan-1, VCAM-1) and the severity-related biomarker (ferritin) directly correlate with expression of inhibitory markers (LILRB1, NKG2A, PD-L1) and inversely correlate with the frequencies of activated and cytotoxic NK-cells (CD69^+^, NKG2D^+^, GzmB^+^ perforin^+^; **Fig.6E**). Plasma levels of soluble MICB, the ligand of NKG2D, correlate inversely with ferritin, suggesting that MICB-NKG2D interactions associate with less severe disease. Accordingly, NKG2D^+^ NK-cell and CD8^+^ T-cells are significantly increased in non-SD compared to SD patients (**Fig.6C** and **Supplementary Fig.S5**). These analyses highlight the association of different dengue severity-related markers with NK-cell impairment.

Unsupervised UMAP/FlowSOM analyses show similar findings (**Fig.6F-H, I-K**). SD patients display increased clusters of NK-cells expressing the inhibitory receptors LILRB1 and NKG2A and non-proliferating NK-cells expressing NKp44, NKp46 and NKG2D and NKG2A (**Fig.6F-H**, respectively clusters 8 and 10). Conversely, activated NK-cells with a more mature phenotype expressing NKG2C and CD57 are present at higher frequencies in non-SD compared to SD patients (**Fig.6F-H,** cluster 1). In a second flow cytometry panel, the key difference between non-SD and SD patients is observed in cluster 8, with a significantly increased percentage in SD patients. These NK-cells are highly activated and proliferating and express high levels of four inhibitory receptors (LAG-3, TIGIT, PD-1, and PD-L1). Clusters 1 and 3 represent activated NK-cells with increased cytotoxic capacity which show a trend of decreased frequency in SD compared to non-SD patients (**Fig.6I-K)**. In LDA analysis, the phenotypic and functional features of NK-cells early in infection (T1) can discriminate patients that develop SD from non-SD patients (**Fig.6L**). NK-cell features are largely overlapping in HW, OW and OB patients (**Fig.6M**) although manual gating analyses revealed decreased GzmB^+^perforin^+^ cells and LILRB1 expression in OW/OB compared to HW non-SD patients (**Supplementary Fig.S6**).

We next determined whether NK-cells from SD patients were altered in their effector function compared to non-SD patients. To this end, NK-cells were assessed for natural cytotoxicity (degranulation assessed as CD107a expression) and production of IFN-γ, TNF-α and MIP-1β after co-culture with K562 target cells with or without cytokine stimulation (IL-12/18). In response to stimulation with K562 cells, NK-cells from SD and non-SD patients produce similar amounts of IFN-γ and no TNF-α or MIP-1β. However, in this condition NK-cell degranulation (CD107a) is significantly decreased in NK-cells from SD patients (**Fig.6N-Q**). IL-12/18 stimulation induces NK-cell production of IFN-γ and boosts all 4 measured functions when combined with K562 stimulation, but it was not able to restore NK-cell degranulation in SD to the levels observed in non-SD patients.

Collectively our data demonstrates phenotypically and functionally altered NK-cells early in infection prior to or at early onset of SD.

### Impaired type-I IFN signalling in SD

We next investigated whether defects in innate viral-sensing could underlie T and NK-cell impairment in SD. To achieve this, we performed gene expression analysis by single-cell RNA-sequencing (scRNA-seq, BD Rhapsody) of PBMCs from 24 dengue patients at T1 (N=12 SD; N=12 non-SD, with each group including 6 HW and 6 OW/OB patients matched by sex and age). scRNA-seq captured gene expression of circulating CD8^+^/CD4^+^ T-cells, NK-cells, MAIT cells, γδ T-cells, B-cells, plasmablasts, monocytes, and dendritic cells (**Fig.7A**). Plasmablast frequency was increased in SD compared to non-SD (**Fig.7B**), consistent with our flow cytometry data (**Supplementary Fig.S8**). Differential gene expression analysis revealed a blunted type-I IFN response in SD compared to non-SD patients in all cell types combined, as well as in CD8^+^ T-cells and NK-cells (**Fig.7C, Supplementary Fig.S7 and Table S2**). Genes significantly downregulated in SD (Bonferroni adjusted p-values) encode proteins driving expression of type-I IFNs or induced by type-I IFNs. These include IFN-induced transmembrane protein (IFITM) 1, 2 and 3, IFN regulatory factor family (IRF) 4, 7 and 9, IFN-stimulated gene 15 (ISG15), IFN-induced helicase C domain-containing protein 1 (IFIH1), IFN inducible protein 16 (IFI16), 2’-5’-oligoadenylate synthetase 1 (OAS1), and signal transducer and activator of transcription (STAT) 1 and 2. IFNAR1 and IFNAR2 as well as IL-2R gamma subunit, which is common to different interleukin receptors, were also significantly downregulated in SD, suggesting decreased responsiveness to cytokines (**Supplementary Table S2**).

**Fig.7.**
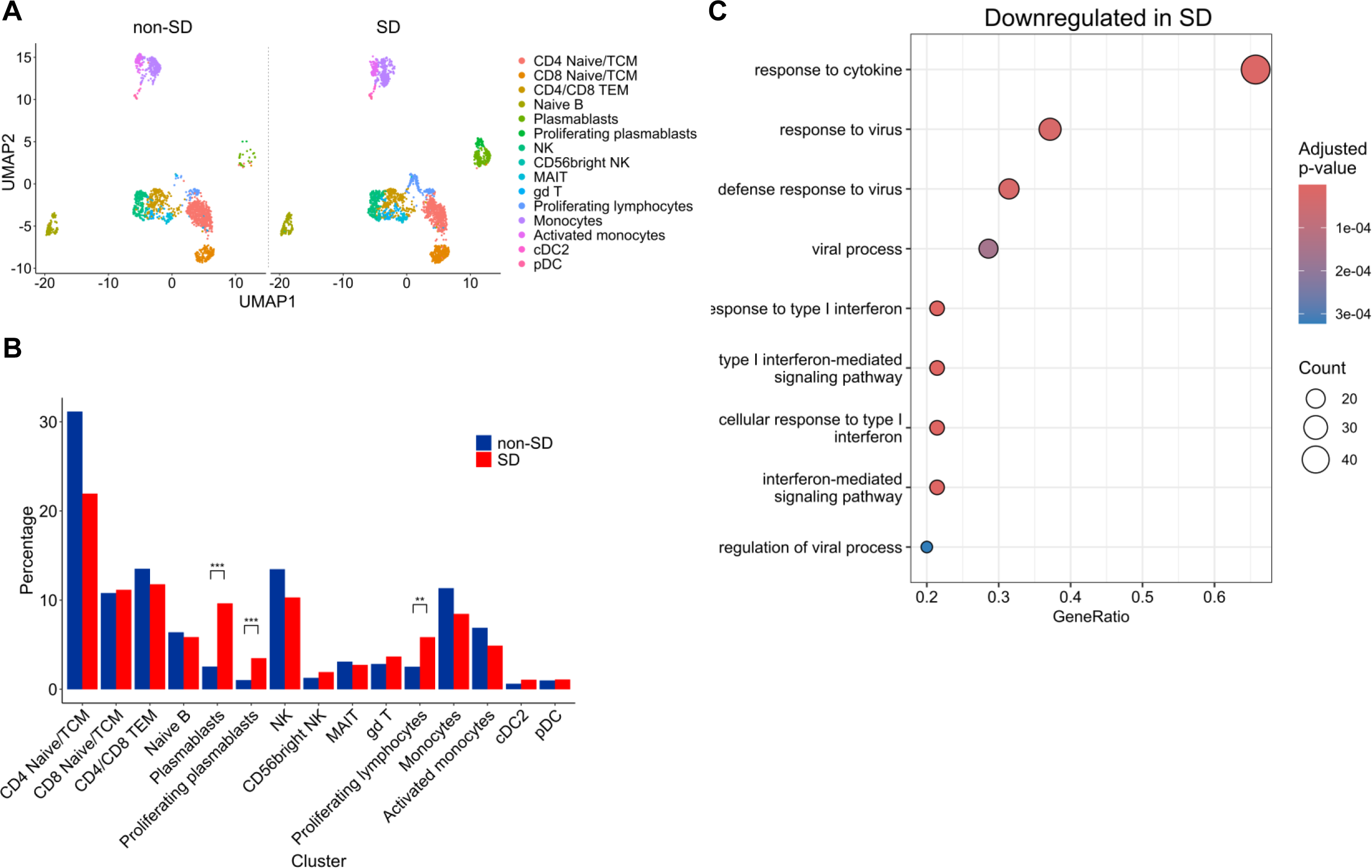
Impaired type-I IFN responses in SD. **(A)** UMAP of scRNA-seq data from PBMCs, coloured by cell type and split by dengue severity. Data from N=24 dengue patients (N=12 SD; N=12 non-SD). **(B)** Percentage of each cell type in non-SD and SD patients. Differential abundance analysis was performed using edgeR, ***p<.001. **(C)** Over-representation analysis of genes significantly downregulated (all cell types combined) in SD vs non-SD. Significant non-redundant gene ontology terms and associated Benjamini-Hochberg adjusted p-values are shown. Count=number of differentially expressed genes (DEGs) in the gene set. GeneRatio=fraction of DEGs in the gene set.

Type-I IFNs mediate NK-cell activation in viral infection, including dengue^36^ and are critical for survival of activated CD8^+^ T-cells^37^. Our data therefore suggests defective virus-sensing and type-I IFN responses as a potential mechanism underlying NK and T-cell dysfunction in SD.

## Discussion

The lack of validated correlates of protection and immunopathology for dengue represents a major challenge for the design of protective vaccines and host-directed therapeutics. Here we provide an in-depth analysis of T and NK-cell signatures associated with dengue disease severity and obesity. Our work provides unparalleled data encompassing phenotypic and functional profiles of T and NK-cells in a unique dengue cohort including a larger number of SD patients than previously analysed in a single study (N=30 SD; N=94 non-SD), as well as sex, age and illness phase matched patients with HW or OW/OB. The latter allows the analysis of the impact of OW/OB, a risk factor for dengue, on immunity to DENV. We show that in early disease CD4^+^/CD8^+^ T and NK-cell profiles linked to activation, proliferation, cytotoxicity, and skin/peripheral tissue-homing, discriminate patients that will progress to SD from those that do not. Within non-SD patients, these profiles could discriminate patients with and without warning signs suggesting that certain warning signs, may be immunologically driven. T/NK-cell profiles were distinct between patients with HW, OW and OB, suggesting BMI impacts immunity to dengue. Furthermore, NK-cell expression of activating/inhibitory receptors and cytotoxic molecules at T1 also clearly discriminates between SD and non-SD patients. Here, the LDA analyses showed a less clear discrimination of patients based on BMI, suggesting NK-cell responses in dengue may be less impacted by host BMI. For all the above T1 analyses we included SD patients enrolled at day 3, prior to development of severe manifestations, as well as SD patients who were recruited in the ICU at day 5, and hence were at early onset of SD. T/NK-cell profiles were similar in SD patients at days 3 and 5 and hence these data were pooled together in our analyses. Our data suggest that T/NK-cells responses: (i) could potentially have prognostic value for early stratification of patients more likely to progress to SD; (ii) are critical for the early anti-viral response and (iii) may represent a novel therapeutic target to restore effective anti-viral immunity in dengue. Each of these points is discussed below.

While most DENV infections are self-limiting, the high number of symptomatic DENV infections during seasonal epidemics rapidly overwhelms health care systems in dengue-endemic countries. The availability of biomarkers for early identification of patients who will develop SD would improve healthcare effectiveness and patient outcomes. Several studies have proposed candidate biomarkers for SD, but these have shown limited clinical value due to their appearance later in disease or their short half-life^38^. A recent study including >7,400 participants proposed a combination of inflammatory (IL-8, CXCL10/IP-10, sTREM-1, and sCD163) and vascular markers (syndecan-1) as potential biomarkers for severe/moderate dengue, all related to excessive activation of macrophages/myeloid cells, the main targets of DENV infection^23^. Excessive macrophage activation is consistent with a scenario of dysfunctional cytotoxic NK/T-cells leading to impaired clearance of virus-infected cells observed in our study.

Our data suggests that suboptimal type-I IFN responses may underlie defective viral clearance and NK/T-cell impairment in SD. Type-I IFN signalling leads to activation of IFN-stimulated genes (ISGs) and induction of an anti-viral state within the cell^39^. Furthermore, type-I IFNs promote NK-cell function and type-I IFN blockade was shown to inhibit NK-cell responses to DENV-infected cells *in vitro*^36^. Similarly, type-I IFNs directly support survival, clonal expansion and cytotoxicity of activated T-cells^37,40,41^. Our findings are consistent with *in vitro* data showing DENV NS5-mediated inhibition of type-I IFN signalling through degradation of STAT2, which was also decreased in SD in our study^42^. Our findings also support early DNA microarray studies showing decreased transcripts of many genes induced by type-I IFN in patients with dengue shock syndrome compared to those without^42–44^.

Antibody dependent enhancement (ADE) of infection mediated by pre-existing subneutralizing DENV-specific IgG antibodies is associated with more severe clinical outcomes^12^. This is mediated by binding of DENV IgG Fc portions to Fc-gamma receptors (FcγRs) on the surface of myeloid cells, leading to DENV internalization and augmented viral replication. Coligation of FcγR and the inhibitory receptor LILRB1 by antibody-opsonized DENV was shown to inhibit FcγR signalling and induction of ISGs. In NK-cells this interaction may result in reduced ability to mediate antibody dependent cellular cytotoxicity, further inhibiting virus clearance^45^. Therefore ADE, decreased type-I IFN signalling, T/NK-cell cytotoxic impairment and decreased survival of DENV-specific T-cells, may all represent interconnected events leading to increased viraemia and severe outcomes. Genetic factors (e.g., MICB, NKG2D SNPs) may further contribute to the suboptimal NK/T-cell response in these patients, as they could render individuals less efficient at mounting cytotoxic NK/T-cells.

Increased viraemia may lead to increased antigen presentation and excessive T-cell activation causing upregulation of T-cell exhaustion markers and increased apoptosis. As SD associates with secondary infections, preferential reactivation of pre-existing, low affinity memory T-cells could also contribute to the altered T-cell response. This is consistent with the higher frequencies of DENV-specific T-cells at T1 in SD versus non-SD. However, our analysis of the capacity of DENV-specific T-cells to recognize NS3 DENV1-4 peptides in secondary DENV2 infected patients did not show evidence of skewing of the T-cell response towards heterologous DENV serotypes. Due to insufficient number of cells at T1, we were only able to test recognition of all 4 serotypes at T2. It therefore remains possible that suboptimal cross-reactive T-cells that are preferentially activated at T1 may have undergone apoptosis and are not detectable at T2, this is in line with their high CD95/Fas expression levels.

Our data is in line with studies in Colombian dengue cohorts showing an NK-cell related signal in SD using bulk RNA-seq^46^ or using virus-inclusive scRNA-seq on a small cohort of 12 non-SD and 7 SD patients^47^. Another study by the same group demonstrated high PD-L1 expression in SD across different cell types, including myeloid cells^22^. These cells could potentially provide ligands for T/NK-cell expressed PD-1. DENV infection was shown to upregulate HLA-class I and non-classical HLA class I molecules such as HLA-E which are ligands for NK-cell inhibitory receptors, respectively LILRB1 and NKG2A, expressed in SD^48^, suggesting that DENV has evolved strategies to counteract the early NK-cell response. It remains to be determined whether DENV proteins may cause upregulation of ligands binding to inhibitory receptors expressed by T-cells to evade the T-cell response.

Lastly, we propose that T/NK-cell impairment may represent a promising therapeutic target for dengue, supporting recent interest of evaluating immune checkpoint blockade in infectious diseases^49^. In acute hepatitis C virus (HCV) infection, PD-1 expression was shown to correlate with CD8^+^ T-cell exhaustion and PD(L)1 blockade could restore the function of these cells^50^. As for HCV patients, higher PD-1 T-cell expression in SD versus non-SD patients does not reflect the higher activation state of these cells, as expression of other activation markers such as HLA-DR, CD38 and CD69 are similar in T-cells from the two patient groups. Previous studies by us and others reported PD-1 expression in memory DENV-specific CD8^+^ T-cells in convalescent patients or healthy donors from a dengue hyperendemic region^51–53^, although these studies did not address links with SD. Here we show that PD-1/PDL1 blockade can restore the cytotoxic potential of CD8^+^ T-cells in some patients. Recent work in a symptomatic *Ifnar*^-/-^ dengue mouse model shows a similar increase of PD-1^+^ CD8^+^ T-cells upon infection with a non-mouse adapted DENV strain which leads to plasma leakage and death. Anti-PD-1 blockade prior to DENV infection significantly improved mouse survival and rescued CD8^+^ T-cell numbers, suggesting a role of PD-1 in T-cell apoptosis^54^. These data in humans and mice support the need for further studies evaluating the impact of checkpoint inhibitors or combinations of checkpoint inhibitors on NK/T-cell function in dengue. Studies in non-human primates show that while viraemia is cleared around day 5 from illness onset similarly to patients, DENV antigens can be detected in tissues until at least day 8^55^. This data supports an important role of cytotoxic NK-cells/DENV-specific T-cells beyond the blood viraemic phase, for clearing reservoirs of DENV-infected cells within tissues.

In summary, our work demonstrates T and NK-cell impairment in SD patients which precedes the development of SD and is present during the critical phase. We propose that these signatures could potentially be used as prognostic markers for SD and represent a novel therapeutic avenue for dengue aimed at restoring their anti-viral function.

## Acknowledgments

The authors wish to acknowledge the assistance of Dr Andrew Herman, Helen Rice, Poppy Miller, Celyn Dugdale and the University of Bristol Faculty of Biomedical Sciences Flow Cytometry Facility. We are grateful to all patients and their families for participating in this study. This study was supported by the Academy of Medical Sciences and the Springboard Award scheme funders: the Wellcome Trust, the Government Department of Business, Energy and Industrial Strategy and the British Heart Foundation and Diabetes UK [SBF007\100173]; the Royal Society (RGS\R1\221078). LG is supported by the Wellcome Trust (222426/Z/21/Z). MS was supported by the Elizabeth Blackwell Institute for Health Research, University of Bristol with funding from the University’s alumni and friends (TRACK award to LR).

## Author contributions

Conceptualization, L.R. and S.Y.; Methodology, L.R., M.G., M.S., C.L., P.K.; Validation, M.G., L.R., M.S., D.D., M.N.; Formal Analysis, M.G., M.S., D.D., L.C.G., N.R., M.N., N.L.V.; Investigation, M.G., M.S., D.D., E.J., N.R., N.M.N., T.T.V., H.Q.C., N.T.X.C., D.T.H.T., D.H.T.L., C.T.T.; Resources, L.R., S.Y.; Data Curation, M.G., L.C.G.; Writing – Original Draft, M.G. and L.R., Writing – Review & Editing, M.G., L.R., S.Y., P.K., L.C.G., N.M.N., H.V.T.M., H.Q.C; Visualization, M.G., L.R., L.C.G., M.N.; Supervision, L.R.; Project Administration, L.R.; Funding Acquisition, L.R. and S.Y.

## Declaration of interests

The authors declare no competing interests.

## STAR Methods

### Ethics approvals

The study protocol, consent and assent forms, and patient information sheets were approved by the ethics committees at the Hospital for Tropical Diseases in Ho Chi Minh City (CS/BND/19/34), the Ministry of Health in Vietnam (24/CN-HÐÐÐ) and the Oxford Tropical Research Ethics Committee (REC) (OxREC reference:36-19).

### Dengue patients

After informed consent/assent, hospitalised patients aged 10-30 were recruited into an observational study designed for dengue patients with overweight/obesity and healthy weight at the Hospital for Tropical Diseases, Ho Chi Minh City, Vietnam. All patients had confirmed dengue and ≤72 hours of fever, except for a proportion of severe patients who were admitted in the intensive care units (ICUs) up to day 5 of fever. Each patient with overweight/obesity was matched 1:1 to a healthy weight patient by age group (10-16; >16-21; >21-26; >26-30), sex, admission ward (general or ICU), and illness phase – febrile (fever days 1-3) or critical (fever days 4-5). The enrolment criteria were selected to minimise confounders due to ageing and comorbidities and to maximise recruitment of severe cases. The exclusion criteria included diabetes, hypertension, cardiovascular disease, signs or symptoms of any other acute infectious disease, undernutrition, and pregnancy.

Definition of BMI groups for paediatric patients (10-19 years) was based on the WHO obesity definition using BMI-for-age^56^; for adult patients (20-30 years) BMI status was defined as follows: individuals with overweight/obesity had a BMI ≥25 kg/m^2^, while HW patients had a BMI ≤22 kg/m^2^ but not less than 17 kg/m^2^. The severity grade classification was recorded for all admitted patients following the WHO 2009 guidelines^3^ and based on plasma leakage (grade 0-2)^57^. PBMC isolation and cryopreservation were performed at hospital enrolment (within 3-5 days of fever) and approx. 3 days later, which largely coincided with discharge (6-9 days of illness).

### Serology

Identification of the DENV serotype of infection and quantification of viraemia (RNAemia) was performed by serotype-specific RT-PCR. The majority of patients were infected with DENV2 serotype (n=86), although some cases of DENV1 (n=22) and DENV4 (n=7) serotypes were also detected. RNAemia was below the detection limit at the time of enrolment in the study for 3 SD and 6 non-SD patients (indicated as NEG; **Table S1)**. Primary/secondary dengue infection was determined based on the ratio of dengue IgM/IgG antibody levels through enzyme-linked immunosorbent assay (ELISA), as described^23^. The majority of patients (n=118) included in the study had a secondary infection; there were five cases of primary infection among the non-SD group. Pro-inflammatory chemokines and cytokines were analysed from plasma samples using a Luminex FLEXMAP 3D® using the Inflammation 20-plex human Procartaplex^TM^ panel. Endothelial, pro-inflammatory, and lipid markers and sMICB were measured by ELISA.

### DENV synthetic peptides

NS3 DENV peptide pools comprised of 15-mer peptides overlapping by 10 amino acids and spanning the sequence of NS3 DENV1-4 (accession umbers: DENV1 – MF314188, DENV2 – NP056776, DENV3 – KY921906, DENV4 – KY921909). NS3 peptide pool 1: N=40, pool 2: N=40, pool 3: N=42. Peptide libraries were designed based on the following virus strains: DENV1 – 2016_Singapore_DENV-1_NPHL, DENV2 – Thailand/16681/84, DENV3 – SG(EHI)D3/23167Y15, and DENV4 – SG(EHI)D4/09291Y16. All peptides were purchased from Mimotopes (Australia) with >80% purity. Peptides were prepared as described previously^27^.

### *Ex vivo* PBMC surface and intracellular staining

PBMCs were thawed in RPMI 10% FBS, washed twice and seeded at 0.8-1×10^6^ cells/well in a 96-well plate. Cells were stained with a viability dye (Zombie Aqua) for 10 min, washed with PBS 1% BSA and antibodies targeting surface markers were added and incubated for 20 min at 4°C. After staining, cells were washed and fixed (BD/eBiosciences). After fixation, the cells were washed three times with perm/wash buffer (BD/eBiosciences) and stained intracellularly with a mixture of antibodies targeting cytokines for 30 min at 4°C. The cells were washed before acquisition using the BD Fortessa X20 flow cytometer. List of antibodies and reagents is shown in Key resources table.

### T-cell stimulation with DENV peptide pools

PBMCs were rapidly thawed in RPMI 10% FBS, washed 2x with PBS 1% BSA and then resuspended in AIM-V 2% AB human serum and rested for 18 h at 37°C. After resting, cells were seeded at 1×10^6^ cells/well in a 96-well plate in AIM-V 2% FBS. Cells were stimulated with or without (DMSO only – negative control) peptide pools from DENV1-4 (all 1µg/mL) or with PMA/ionomycin (positive control) for 6 h at 37°C in the presence of brefeldin A (2 µg/mL). To assess degranulation, anti-CD107a antibody was added at the beginning of stimulation. After stimulation, cells were washed and stained as described above. List of antibodies and reagents is shown in Key resources table.

### PD-1/PD-L1 blockade

PBMCs were rapidly thawed, washed twice and then resuspended in AIM-V 2% AB human serum and plated at 0.8-1×10^6^ cells/well in a 96-well plate. Blocking antibodies (αPD-1/PD-L1) or isotype controls (IgG_4_/IgG1) were added at a final concentration of 10 µg/mL. After 2 h pre-incubation, cells were stimulated with NS3 DENV peptide pools (all 1 µg/mL) or with αCD3/CD28 Dynabeads (positive control) for 16 h at 37°C in the presence of brefeldin A (2 µg/mL), anti-CD107a antibody and blocking antibodies. After incubation, cells were washed and stained as described above.

### NK-cell killing assay

PBMCs were thawed, washed with RPMI 10% FBS and rested or stimulated overnight with 10 ng/mL IL-12 and 100 ng/mL IL-18 in RPMI 10% FBS at 37°C. After incubation, K562 cells were added to rested or stimulated PBMCs at an effector to target ratio of 10:1 (1×10^6^ PBMCs: 1×10^6^ K562 cells). Wells containing only PBMCs (no K562 cells) were also included as a negative control. The plate was incubated for 6 h at 37°C in the presence of anti-CD107a, while Monensin (2 µM) and brefeldin A (2 µg/mL) were added 1 hour into the assay. After the incubation, PBMCs were subsequently stained as previously described. List of antibodies is shown in Key resources table.

### BD Rhapsody scRNA-seq library generation and sequencing

The PBMCs were stained with CD45-PE for 30 min at room temperature. Cells were washed twice in FACS buffer (PBS 2% FBS), incubated with Fc block solution for 15 min at room temperature, then each sample labelled with a separate oligonucleotide-conjugated Sample Tag (BD Flex Single-Cell Multiplexing Kits A-D) for 60 min at 4°C. Cells were washed twice in BD Pharmingen Stain buffer. Twelve samples (3500 cells/sample; n=3 from each group - non-SD HW, non-SD OW/OB, SD HW, SD OW/OB) were pooled and loaded onto one BD Rhapsody Cartridge following manufacturer’s instructions. The remaining 12 samples were pooled and loaded onto a second BD Rhapsody Cartridge. Targeted mRNA and Sample Tag library preparation was performed according to manufacturer’s instructions. Targeted mRNA libraries were generated using the BD Rhapsody Immune Response Panel HS and a custom panel containing and additional 145 genes (**Supplementary Table S3**). Libraries were quantified using a Qubit Fluorometer with the Qubit dsDNA HS Assay Kit and their fragment distribution analysed using a TapeStation with the High Sensitivity D5000 ScreenTape Asay (Agilent). Sequencing was performed on an Illumina NovaSeq X Plus (PE150).

### BD Rhapsody scRNA-seq analysis

BCL files were converted to FASTQ files using Illumina bcl2fastq. FASTQ files were processed to a gene-by-cell matrix (one per BD Rhapsody Cartridge) using the BD Rhapsody Sequence Analysis Pipeline (v2.2.1). Downstream analysis was performed in R (v4.4.0). A Seurat^59^ (v5.1.0) object was generated from distribution-based error correction-adjusted molecule counts (two Cartridges combined) and filtered to retain cells with a defined Sample Tag and to remove doublets (cell labels associated with more than one Sample Tag). Cells with low unique molecular identifier counts (<125) and/or low gene counts (<40) were removed. Count data was analysed following the standard Seurat pipeline. In brief, counts were normalised by library size, scaled, and principal component analysis performed using the top 200 variable genes. A shared nearest neighbour graph was constructed using the top 20 principal components and Louvain clusters identified (resolution 0.5). UMAP was performed using top 20 principal components. Reference-based cell annotation was performed using Azimuth^60^ (v0.5.0) with the PBMC reference dataset. Cluster markers were identified using the Wilcoxon rank sum test (Seurat FindAllMarkers function, only.pos=TRUE) and clusters assigned to cell types using expert knowledge. Final cell annotations were determined using a combination of Azimuth results and manual cluster assignments.

Differential gene expression analysis between non-SD and SD (HW and OW/OB patients combined) was performed for all cell types combined, for total CD8^+^ T-cells, or for total NK cells, using the Wilcoxon rank sum test (Seurat FindMarkers Function, logfc.threshold=0). Over-representation analysis for Gene Ontology Biological Process terms was performed using cluster Profiler^61^ (v4.12.2) for genes significantly downregulated in SD compared with non-SD. Redundant Gene Ontology terms were removed using the clusterProfiler simplify function. Differential cell type abundance analysis was performed using edgeR^62,63^ (v4.2.1). Namely, a common dispersion values across all clusters was estimated using the estimateDisp function (trend.method=”"none”). Quasi-likelihood negative binomial generalised log-linear models were fit to per sample cluster counts (glmQLFit function, robust=TRUE, abundance.trend=FALSE) and quasi F-tests (qlmQLFTest function) employed to test for differential abundance between non-SD and SD (HW and OW/OB patients combined).

### Data and statistical analyses

Analysis of flow cytometry data, including compensation was performed using FlowJo v10.10.0. Quality control was performed using the FlowJo plugin – PeacoQC. Statistical analysis and data visualisation were performed using R v4.2.1. Normality of data distribution was assessed by Shapiro-Wilk test. Differences between two patient groups were calculated by Mann-Whitney U test, for more than two groups we used one-way ANOVA (Kruskal-Wallis) followed by Dunn’s test for multiple comparisons adjusted using the Benjamini-Hochberg method to control for FDR. Analysis between paired samples was performed using Wilcoxon test. Correlation analysis was performed using Spearman’s rank correlation test with/without correction for multiple comparisons. P-values are indicated as follows: *p<.05, **p<.01, ***p<.001, ****p<.0001. The LDA analysis combined clinical information and frequencies/MFIs of markers in the cell subsets, and it was calculated using MASS package. Data analyses was performed by grouping data by specific time points (T1 and T2) as well as by day of fever - we report the most informative analysis.

## Supplemental information

**Fig.S1. Distinct T and NK-cell profiles associate with dengue severity.** Linear discriminant analysis separating dengue patients at **(A,C)** T1 (N=42) and **(B,D)** T2 (N=84). Data points represent individual patients from dengue, dengue with warning signs, and severe dengue group (colour) and based on the **(A,B)** BMI status - healthy weight, overweight, and obesity or **(C,D)** severity status – admitted and progressed group (shape). Ellipses represent 95% confidence intervals. LD1 and LD2 were derived using all features shown in Fig.1D-I.

**Fig.S2. DENV NS3-specific T-cell response.** Single cytokine response by **(A)** CD4^+^ and **(B)** CD8^+^ T-cells in non-SD and SD patients at T2 (N=37) following NS3 DENV1-4 peptide stimulation.

**Fig.S3. Assessment of DENV2-specific T-cell function during DENV2 infection.** Pie charts showing the number of functions simultaneously exhibited by **(A)** CD4^+^ and **(B)** CD8^+^ T-cells following NS3 DENV2 peptide stimulation. The different shades of grey represent the range of 1-5 functions, the outer arcs indicate the specific functions (IFN-γ/TNF-α/IL-2/MIP-1β/CD107a) defined by Boolean gating.

**Fig.S4. Metabolic activity measured by SCENITH. (A)** Fatty acid and amino acid oxidation (FAO&AAO) capacity, **(B)** glycolytic capacity, **(C)** mitochondrial dependence, and **(D)** glucose dependence. SCENITH was performed and capacities and dependencies were calculated according to Arguello et al.^32^ and Luscombe et al.^33^.

**Fig.S5. Impaired phenotype of CD8^+^ T-cells in SD.** Frequency of total or Ki-67^+^ CD8^+^ T-cells expressing NKG2D in non-SD and SD patient group.

**Fig.S6. Impaired phenotype of Ki-67^+^ total NK cells in SD.** Expression of **(A)** activating and **(B)** inhibitory receptors by Ki-67^+^ total NK-cells in HW (green) and OW/OB (orange) non-SD and SD patients.

**Fig.S7. Impaired type-I IFN responses in SD.** Over-representation analysis of genes significantly downregulated in SD vs non-SD in CD8^+^ T-cells **(A)** and NK-cells **(B)**. All significant or top 15 non-redundant gene ontology terms and associated Benjamini-Hochberg adjusted p-values are shown. Count=number of differentially expressed genes (DEGs) in the gene set. GeneRatio=fraction of DEGs in the gene set. Data from N=24 dengue patients (N=12 non-SD; N=12 SD).

**Fig.S8. Increased frequency of B-cells and plasmablasts in SD. (A)** Frequency of CD3^-^ CD19^+^ **(B)** Gating strategy and frequency of plasmablasts in non-SD and SD patients [HW (green) and OW/OB (orange)]. Data from N=94 dengue patients at T2 (N=68 non-SD; N=26 SD).

**Table S1.** Summary table of clinical information at the admission time point

**Table S2.** List of downregulated genes (all cell types combined) in SD patient group.

**Table S3.** List of custom genes from BD Rhapsody gene panel.

## Notes

### Competing Interest Statement

The authors have declared no competing interest.

## References

1. Bhatt, S., Gething, P.W., Brady, O.J., Messina, J.P., Farlow, A.W., Moyes, C.L., Drake, J.M., Brownstein, J.S., Hoen, A.G., Sankoh, O., et al. (2013). The global distribution and burden of dengue. Nature 496, 504–507. 10.1038/nature12060.

2. Yacoub, S., Kotit, S., and Yacoub, M.H. (2011). Disease appearance and evolution against a background of climate change and reduced resources. Philos. Transact. A Math. Phys. Eng. Sci. 369, 1719–1729. 10.1098/rsta.2011.0013.

3. Dengue: Guidelines for Diagnosis, Treatment, Prevention and Control: New Edition (2009). (World Health Organization).

4. Thomas, S.J. (2023). Is new dengue vaccine efficacy data a relief or cause for concern? Npj Vaccines 8, 1–6. 10.1038/s41541-023-00658-2.

5. Screaton, G., Mongkolsapaya, J., Yacoub, S., and Roberts, C. (2015). New insights into the immunopathology and control of dengue virus infection. Nat. Rev. Immunol. 15, 745–759. 10.1038/nri3916.

6. Trang, N.T.H., Long, N.P., Hue, T.T.M., Hung, L.P., Trung, T.D., Dinh, D.N., Luan, N.T., Huy, N.T., and Hirayama, K. (2016). Association between nutritional status and dengue infection: a systematic review and meta-analysis. BMC Infect. Dis. 16, 172. 10.1186/s12879-016-1498-y.

7. Gallagher, P., Rong, K., Rivino, L., and Yacoub, S. (2020). The association of obesity and severe dengue: possible pathophysiological mechanisms. J. Infect. 81, 10–16. 10.1016/j.jinf.2020.04.039.

8. Rebeles, J., Green, W.D., Alwarawrah, Y., Nichols, A.G., Eisner, W., Danzaki, K., MacIver, N.J., and Beck, M.A. (2019). Obesity-induced changes in T-cell metabolism are associated with impaired memory T-cell response to influenza and are not reversed with weight loss. J. Infect. Dis. 219, 1652–1661. 10.1093/infdis/jiy700.

9. Sheridan, P.A., Paich, H.A., Handy, J., Karlsson, E.A., Hudgens, M.G., Sammon, A.B., Holland, L.A., Weir, S., Noah, T.L., and Beck, M.A. (2012). Obesity is associated with impaired immune response to influenza vaccination in humans. Int. J. Obes. 36, 1072–1077. 10.1038/ijo.2011.208.

10. Tobin, L.M., Hogan, A.E., Shea, D.O., Tobin, L.M., Mavinkurve, M., Carolan, E., Kinlen, D., Brien, E.C.O., Little, M.A., Finlay, D.K., et al. (2017). NK cells in childhood obesity are activated, metabolically stressed, and functionally deficient. JCI Insight 2.

11. Michelet, X., Dyck, L., Hogan, A., Loftus, R.M., Duquette, D., Wei, K., Beyaz, S., Tavakkoli, A., Foley, C., Donnelly, R., et al. (2018). Metabolic reprogramming of natural killer cells in obesity limits antitumor responses. Nat. Immunol. 19, 1330–1340. 10.1038/s41590-018-0251-7.

12. Katzelnick, L.C., Gresh, L., Halloran, M.E., Mercado, J.C., Kuan, G., Gordon, A., Balmaseda, A., and Harris, E. (2017). Antibody-dependent enhancement of severe dengue disease in humans. Science 358, 929–932. 10.1126/science.aan6836.

13. Weiskopf, D., Angelo, M.A., De Azeredo, E.L., Sidney, J., Greenbaum, J.A., Fernando, A.N., Broadwater, A., Kolla, R.V., De Silva, A.D., De Silva, A.M., et al. (2013). Comprehensive analysis of dengue virus-specific responses supports an HLA-linked protective role for CD8+ T cells. Proc. Natl. Acad. Sci. U. S. A. 110. 10.1073/pnas.1305227110.

14. Weiskopf, D., Bangs, D.J., Sidney, J., Kolla, R.V., De Silva, A.D., De Silva, A.M., Crotty, S., Peters, B., and Sette, A. (2015). Dengue virus infection elicits highly polarized CX3CR1+ cytotoxic CD4+ T cells associated with protective immunity. Proc. Natl. Acad. Sci. U. S. A. 112, E4256–E4263. 10.1073/pnas.1505956112.

15. Mongkolsapaya, J., Dejnirattisai, W., Xu, X.N., Vasanawathana, S., Tangthawornchaikul, N., Chairunsri, A., Sawasdivorn, S., Duangchinda, T., Dong, T., Rowland-Jones, S., et al. (2003). Original antigenic sin and apoptosis in the pathogenesis of dengue hemorrhagic fever. Nat. Med. 9, 921– 927. 10.1038/nm887.

16. Duangchinda, T., Dejnirattisai, W., Vasanawathana, S., Limpitikul, W., Tangthawornchaikul, N., Malasit, P., Mongkolsapaya, J., and Screaton, G. (2010). Immunodominant T-cell responses to dengue virus NS3 are associated with DHF. Proc. Natl. Acad. Sci. U. S. A. 107, 16922–16927. 10.1073/pnas.1010867107.

17. Rivino, L. (2016). T cell immunity to dengue virus and implications for vaccine design. Expert Rev. Vaccines 15, 443–453. 10.1586/14760584.2016.1116948.

18. Mangada, M.M., and Rothman, A.L. (2005). Altered Cytokine Responses of Dengue-Specific CD4+ T Cells to Heterologous Serotypes 1. J. Immunol. 175, 2676–2683. 10.4049/jimmunol.175.4.2676.

19. Hatch, S., Endy, T.P., Thomas, S., Mathew, A., Potts, J., Pazoles, P., Libraty, D.H., Gibbons, R., and Rothman, A.L. (2011). Intracellular Cytokine Production by Dengue Virus–specific T cells Correlates with Subclinical Secondary Infection. J. Infect. Dis. 203, 1282–1291. 10.1093/infdis/jir012.

20. Vuong, N.L., Cheung, K.W., Periaswamy, B., Vi, T.T., Duyen, H.T.L., Leong, Y.S., Binte Hamis, Z.N., Gregorova, M., Ooi, E.E., Sessions, O., et al. (2022). Hyperinflammatory Syndrome, Natural Killer Cell Function, and Genetic Polymorphisms in the Pathogenesis of Severe Dengue. J. Infect. Dis. 226, 1338–1347. 10.1093/infdis/jiac093.

21. Shabrish, S., Karnik, N., Gupta, V., Bhate, P., and Madkaikar, M. (2020). Impaired NK cell activation during acute dengue virus infection: A contributing factor to disease severity. Heliyon 6, e04320. 10.1016/j.heliyon.2020.e04320.

22. Robinson, M.L., Glass, D.R., Duran, V., Agudelo Rojas, O.L., Sanz, A.M., Consuegra, M., Sahoo, M.K., Hartmann, F.J., Bosse, M., Gelvez, R.M., et al. (2023). Magnitude and kinetics of the human immune cell response associated with severe dengue progression by single-cell proteomics. Sci. Adv. 9, eade7702. 10.1126/sciadv.ade7702.

23. Vuong, N.L., Lam, P.K., Ming, D.K.Y., Duyen, H.T.L., Nguyen, N.M., Tam, D.T.H., Duong Thi Hue, K., Chau, N.V., Chanpheaktra, N., Lum, L.C.S., et al. (2021). Combination of inflammatory and vascular markers in the febrile phase of dengue is associated with more severe outcomes. eLife 10, e67460. 10.7554/eLife.67460.

24. Libraty, D.H., Zhang, L., Obcena, A.M., Brion, J.D., and Capeding, R.Z. (2014). Circulating levels of soluble MICB in infants with symptomatic primary dengue virus infections. PLoS ONE 9. 10.1371/journal.pone.0098509.

25. Khor, C.C., Chau, T.N.B., Pang, J., Davila, S., Long, H.T., Ong, R.T.H., Dunstan, S.J., Wills, B., Farrar, J., Van Tram, T., et al. (2011). Genome-wide association study identifies susceptibility loci for dengue shock syndrome at MICB and PLCE1. Nat. Genet. 43, 1139–1141. 10.1038/ng.960.

26. Whitehorn, J., Chau, T.N.B., Nguyet, N.M., Kien, D.T.H., Quyen, N.T.H., Trung, D.T., Pang, J., Wills, B., Van Vinh Chau, N., Farrar, J., et al. (2013). Genetic Variants of MICB and PLCE1 and Associations with Non-Severe Dengue. PLoS ONE 8, e59067. 10.1371/journal.pone.0059067.

27. Rivino, L., Kumaran, E.A., Thein, T.L., Too, C.T., Gan, V.C.H., Hanson, B.J., Wilder-Smith, A., Bertoletti, A., Gascoigne, N.R.J., Lye, D.C., et al. (2015). Virus-specific T lymphocytes home to the skin during natural dengue infection. Sci. Transl. Med. 7. 10.1126/scitranslmed.aaa0526.

28. Rivino, L., and Lim, M.Q. (2017). CD4+ and CD8+ T-cell immunity to Dengue – lessons for the study of Zika virus. Immunology 150, 146–154. 10.1111/imm.12681.

29. Rivino, L., Kumaran, E.A.P., Jovanovic, V., Nadua, K., Teo, E.W., Pang, S.W., Teo, G.H., Gan, V.C.H., Lye, D.C., Leo, Y.S., et al. (2013). Differential Targeting of Viral Components by CD4+ versus CD8+ T Lymphocytes in Dengue Virus Infection. J. Virol. 87, 2693–2706. 10.1128/jvi.02675-12.

30. Chandele, A., Sewatanon, J., Gunisetty, S., Singla, M., Onlamoon, N., Akondy, R.S., Kissick, H.T., Nayak, K., Reddy, E.S., Kalam, H., et al. (2016). Characterization of Human CD8 T Cell Responses in Dengue Virus-Infected Patients from India. J. Virol. 90, 11259–11278. 10.1128/JVI.01424-16.

31. Wherry, E.J. (2011). T cell exhaustion. Nat. Immunol. 12, 492–499. 10.1038/ni.2035.

32. Argüello, R.J., Combes, A.J., Char, R., Gigan, J.P., Baaziz, A.I., Bousiquot, E., Camosseto, V., Samad, B., Tsui, J., Yan, P., et al. (2020). SCENITH: A Flow Cytometry-Based Method to Functionally Profile Energy Metabolism with Single-Cell Resolution. Cell Metab. 32, 1063–1075.e7. 10.1016/j.cmet.2020.11.007.

33. Luscombe, C., Jones, E., Gregorova, M., Jones, N., and Rivino, L. (2024). Impact of cryopreservation on immune cell metabolism as measured by SCENITH. Preprint at bioRxiv, 10.1101/2024.06.12.598758 https://doi.org/10.1101/2024.06.12.598758.

34. Lanier, L.L. (1998). NK CELL RECEPTORS. Annu. Rev. Immunol. 16, 359–393. 10.1146/annurev.immunol.16.1.359.

35. Zimmer, C.L., Cornillet, M., Solà-Riera, C., Cheung, K.W., Ivarsson, M.A., Lim, M.Q., Marquardt, N., Leo, Y.S., Lye, D.C., Klingström, J., et al. (2019). NK cells are activated and primed for skin-homing during acute dengue virus infection in humans. Nat. Commun. 10. 10.1038/s41467-019-11878-3.

36. Lim, D.S.L., Yawata, N., Selva, K.J., Li, N., Tsai, C.Y., Yeong, L.H., Liong, K.H., Ooi, E.E., Chong, M.K., Ng, M.L., et al. (2014). The Combination of Type I IFN, TNF-α, and Cell Surface Receptor Engagement with Dendritic Cells Enables NK Cells To Overcome Immune Evasion by Dengue Virus. J. Immunol. 193, 5065–5075. 10.4049/jimmunol.1302240.

37. Marrack, P., Kappler, J., and Mitchell, T. (1999). Type I Interferons Keep Activated T Cells Alive. J. Exp. Med. 189, 521–530. 10.1084/jem.189.3.521.

38. Rathore, A.P., Farouk, F.S., and St. John, A.L. (2020). Risk factors and biomarkers of severe dengue. Curr. Opin. Virol. 43, 1–8. 10.1016/j.coviro.2020.06.008.

39. McNab, F., Mayer-Barber, K., Sher, A., Wack, A., and O’Garra, A. (2015). Type I interferons in infectious disease. Nat. Rev. Immunol. 15, 87–103. 10.1038/nri3787.

40. Kolumam, G.A., Thomas, S., Thompson, L.J., Sprent, J., and Murali-Krishna, K. (2005). Type I interferons act directly on CD8 T cells to allow clonal expansion and memory formation in response to viral infection. J. Exp. Med. 202, 637–650. 10.1084/jem.20050821.

41. Kohlmeier, J.E., Cookenham, T., Roberts, A.D., Miller, S.C., and Woodland, D.L. (2010). Type I interferons regulate cytolytic activity of memory CD8+ T cells in the lung airways during respiratory virus challenge. Immunity 33, 96–105. 10.1016/j.immuni.2010.06.016.

42. Morrison, J., Laurent-Rolle, M., Maestre, A.M., Rajsbaum, R., Pisanelli, G., Simon, V., Mulder, L.C.F., Fernandez-Sesma, A., and García-Sastre, A. (2013). Dengue Virus Co-opts UBR4 to Degrade STAT2 and Antagonize Type I Interferon Signaling. PLOS Pathog. 9, e1003265. 10.1371/journal.ppat.1003265.

43. Simmons, C.P., Popper, S., Dolocek, C., Chau, T.N.B., Griffiths, M., Dung, N.T.P., Long, T.H., Hoang, D.M., Chau, N.V., Thao, L.T.T., et al. (2007). Patterns of Host Genome—Wide Gene Transcript Abundance in the Peripheral Blood of Patients with Acute Dengue Hemorrhagic Fever. J. Infect. Dis. 195, 1097–1107. 10.1086/512162.

44. Ashour, J., Laurent-Rolle, M., Shi, P.-Y., and García-Sastre, A. (2009). NS5 of Dengue Virus Mediates STAT2 Binding and Degradation. J. Virol. 83, 5408–5418. 10.1128/jvi.02188-08.

45. Chan, K.R., Ong, E.Z., Tan, H.C., Zhang, S.L.-X., Zhang, Q., Tang, K.F., Kaliaperumal, N., Lim, A.P.C., Hibberd, M.L., Chan, S.H., et al. (2014). Leukocyte immunoglobulin-like receptor B1 is critical for antibody-dependent dengue. Proc. Natl. Acad. Sci. 111, 2722–2727. 10.1073/pnas.1317454111.

46. Robinson, M., Sweeney, T.E., Barouch-Bentov, R., Sahoo, M.K., Kalesinskas, L., Vallania, F., Sanz, A.M., Ortiz-Lasso, E., Albornoz, L.L., Rosso, F., et al. (2019). A 20-Gene Set Predictive of Progression to Severe Dengue. Cell Rep. 26, 1104–1111.e4. 10.1016/j.celrep.2019.01.033.

47. Ghita, L., Yao, Z., Xie, Y., Duran, V., Cagirici, H.B., Samir, J., Osman, I., Rebellón-Sánchez, D.E., Agudelo-Rojas, O.L., Sanz, A.M., et al. (2023). Global and cell type-specific immunological hallmarks of severe dengue progression identified via a systems immunology approach. Nat. Immunol. 24, 2150–2163. 10.1038/s41590-023-01654-3.

48. McKechnie, J.L., Beltrán, D., Pitti, A., Saenz, L., Araúz, A.B., Vergara, R., Harris, E., Lanier, L.L., Blish, C.A., and López-Vergès, S. (2019). HLA Upregulation During Dengue Virus Infection Suppresses the Natural Killer Cell Response. Front. Cell. Infect. Microbiol. 9, 268. 10.3389/fcimb.2019.00268.

49. Wykes, M.N., and Lewin, S.R. (2018). Immune checkpoint blockade in infectious diseases. Nat. Rev. Immunol. 18, 91–104. 10.1038/nri.2017.112.

50. Urbani, S., Amadei, B., Tola, D., Massari, M., Schivazappa, S., Missale, G., and Ferrari, C. (2006). PD-1 Expression in Acute Hepatitis C Virus (HCV) Infection Is Associated with HCV-Specific CD8 Exhaustion. J. Virol. 80, 11398–11403. 10.1128/jvi.01177-06.

51. de Alwis, R., Bangs, D.J., Angelo, M.A., Cerpas, C., Fernando, A., Sidney, J., Peters, B., Gresh, L., Balmaseda, A., de Silva, A.D., et al. (2016). Immunodominant Dengue Virus-Specific CD8+ T Cell Responses Are Associated with a Memory PD-1+ Phenotype. J. Virol. 90, 4771–4779. 10.1128/JVI.02892-15.

52. Chng, M.H.Y., Lim, M.Q., Rouers, A., Becht, E., Lee, B., MacAry, P.A., Lye, D.C., Leo, Y.S., Chen, J., Fink, K., et al. (2019). Large-Scale HLA Tetramer Tracking of T Cells during Dengue Infection Reveals Broad Acute Activation and Differentiation into Two Memory Cell Fates. Immunity 51, 1119–1135.e5. 10.1016/j.immuni.2019.10.007.

53. Rouers, A., Chng, M.H.Y., Lee, B., Rajapakse, M.P., Kaur, K., Toh, Y.X., Sathiakumar, D., Loy, T., Thein, T.-L., Lim, V.W.X., et al. (2021). Immune cell phenotypes associated with disease severity and long-term neutralizing antibody titers after natural dengue virus infection. Cell Rep. Med. 2, 100278. 10.1016/j.xcrm.2021.100278.

54. Idris, F., Ting, D.H.R., Tan, E.T.X., Wan, C., Chan, K.R., Benke, P.I., Marzinek, J.K., Copping, J.M., Li, Q.H., Walsh, I., et al. (2024). De-glycosylated non-structural protein 1 enhances dengue virus clearance by limiting PD-L1/PD-1 mediated T cell apoptosis. Preprint at bioRxiv, 10.1101/2024.01.08.574590 https://doi.org/10.1101/2024.01.08.574590.

55. Marchette, N.J., Halstead, S.B., Falkler, W.A., Jr., Stenhouse, A., and Nash, D. (1973). Studies on the Pathogenesis of Dengue Infection in Monkeys. III. Sequential Distribution of Virus in Primary and Heterologous Infections. J. Infect. Dis. 128, 23–30. 10.1093/infdis/128.1.23.

56. Obesity and overweight https://www.who.int/news-room/fact-sheets/detail/obesity-and-overweight.

57. Lam, P.K., McBride, A., Le, D.H.T., Huynh, T.T., Vink, H., Wills, B., and Yacoub, S. (2020). Visual and Biochemical Evidence of Glycocalyx Disruption in Human Dengue Infection, and Association With Plasma Leakage Severity. Front. Med. 7.

58. Tran, V.T., Inward, R.P.D., Gutierrez, B., Nguyen, N.M., Nguyen, P.T., Rajendiran, I., Cao, T.T., Duong, K.T.H., Kraemer, M.U.G., and Yacoub, S. (2023). Reemergence of Cosmopolitan Genotype Dengue Virus Serotype 2, Southern Vietnam. Emerg. Infect. Dis. 29, 2180–2182. 10.3201/eid2910.230529.

59. Hao, Y., Stuart, T., Kowalski, M.H., Choudhary, S., Hoffman, P., Hartman, A., Srivastava, A., Molla, G., Madad, S., Fernandez-Granda, C., et al. (2024). Dictionary learning for integrative, multimodal and scalable single-cell analysis. Nat. Biotechnol. 42, 293–304. 10.1038/s41587-023-01767-y.

60. Hao, Y., Hao, S., Andersen-Nissen, E., Mauck, W.M., Zheng, S., Butler, A., Lee, M.J., Wilk, A.J., Darby, C., Zager, M., et al. (2021). Integrated analysis of multimodal single-cell data. Cell 184, 3573–3587.e29. 10.1016/j.cell.2021.04.048.

61. Wu, T., Hu, E., Xu, S., Chen, M., Guo, P., Dai, Z., Feng, T., Zhou, L., Tang, W., Zhan, L., et al. (2021). clusterProfiler 4.0: A universal enrichment tool for interpreting omics data. The Innovation 2, 100141. 10.1016/j.xinn.2021.100141.

62. Chen, Y., Chen, L., Lun, A.T.L., Baldoni, P.L., and Smyth, G.K. (2024). edgeR 4.0: powerful differential analysis of sequencing data with expanded functionality and improved support for small counts and larger datasets. Preprint at bioRxiv, 10.1101/2024.01.21.576131 https://doi.org/10.1101/2024.01.21.576131.

63. Amezquita, R.A., Lun, A.T.L., Becht, E., Carey, V.J., Carpp, L.N., Geistlinger, L., Marini, F., Rue-Albrecht, K., Risso, D., Soneson, C., et al. (2020). Orchestrating single-cell analysis with Bioconductor. Nat. Methods 17, 137–145. 10.1038/s41592-019-0654-x.

