## Supplemental Data for "Early NK-cell and T-cell dysfunction marks progression to severe dengue in patients with obesity and healthy weight"

### Supplementary Figure S1

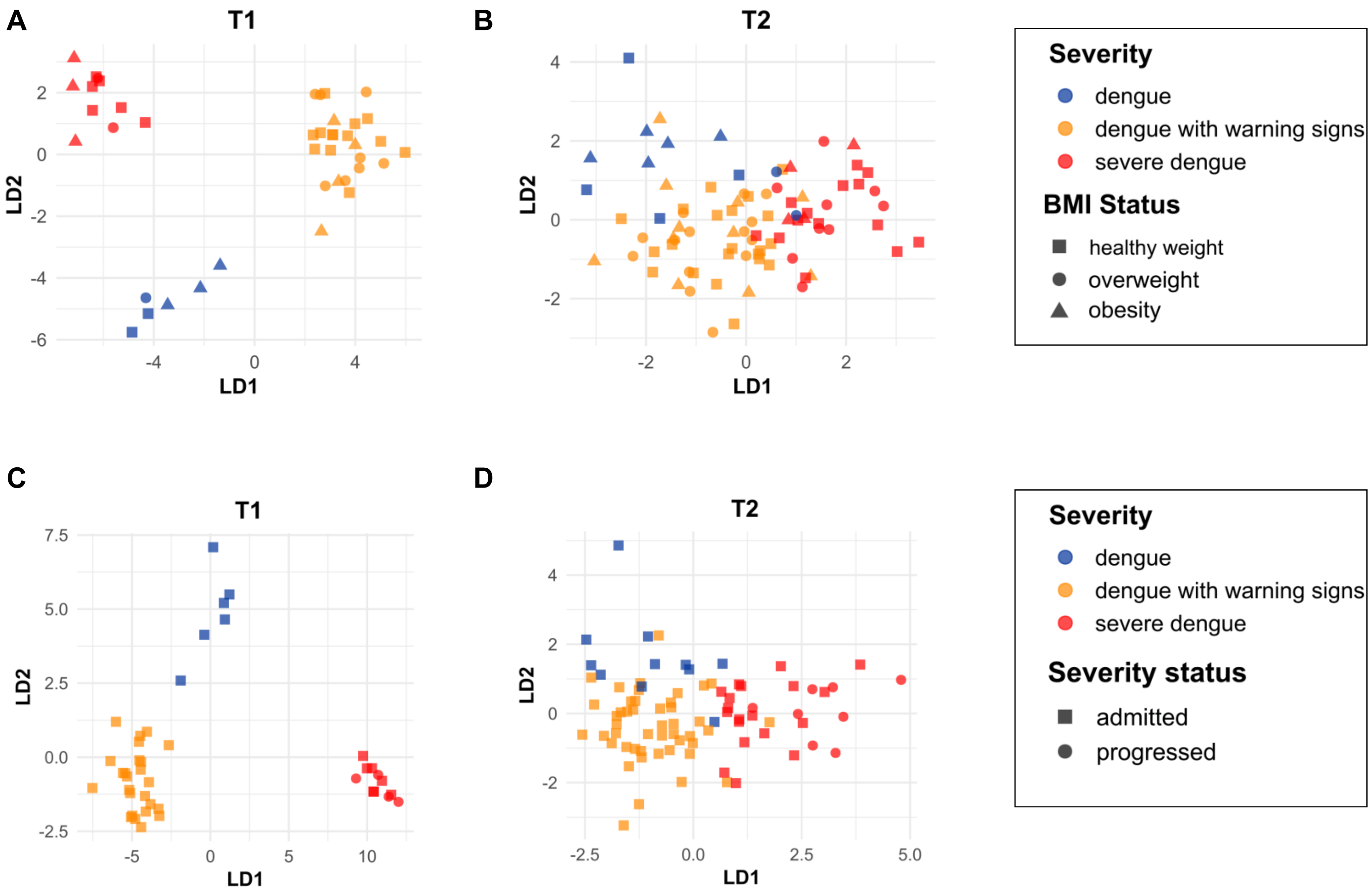


**Fig.S1. Distinct T and NK-cell profiles associate with dengue severity.** Linear discriminant analysis separating dengue patients at **(A,C)** T1 (N=42) and **(B,D)** T2 (N=84). Data points represent individual patients from dengue, dengue with warning signs, and severe dengue group (colour) and based on the **(A,B)** BMI status - healthy weight, overweight, and obesity or **(C,D)** severity status – admitted and progressed group (shape). Ellipses represent 95% confidence intervals. LD1 and LD2 were derived using all features shown in Fig.1D-I.

### Supplementary Figure S2

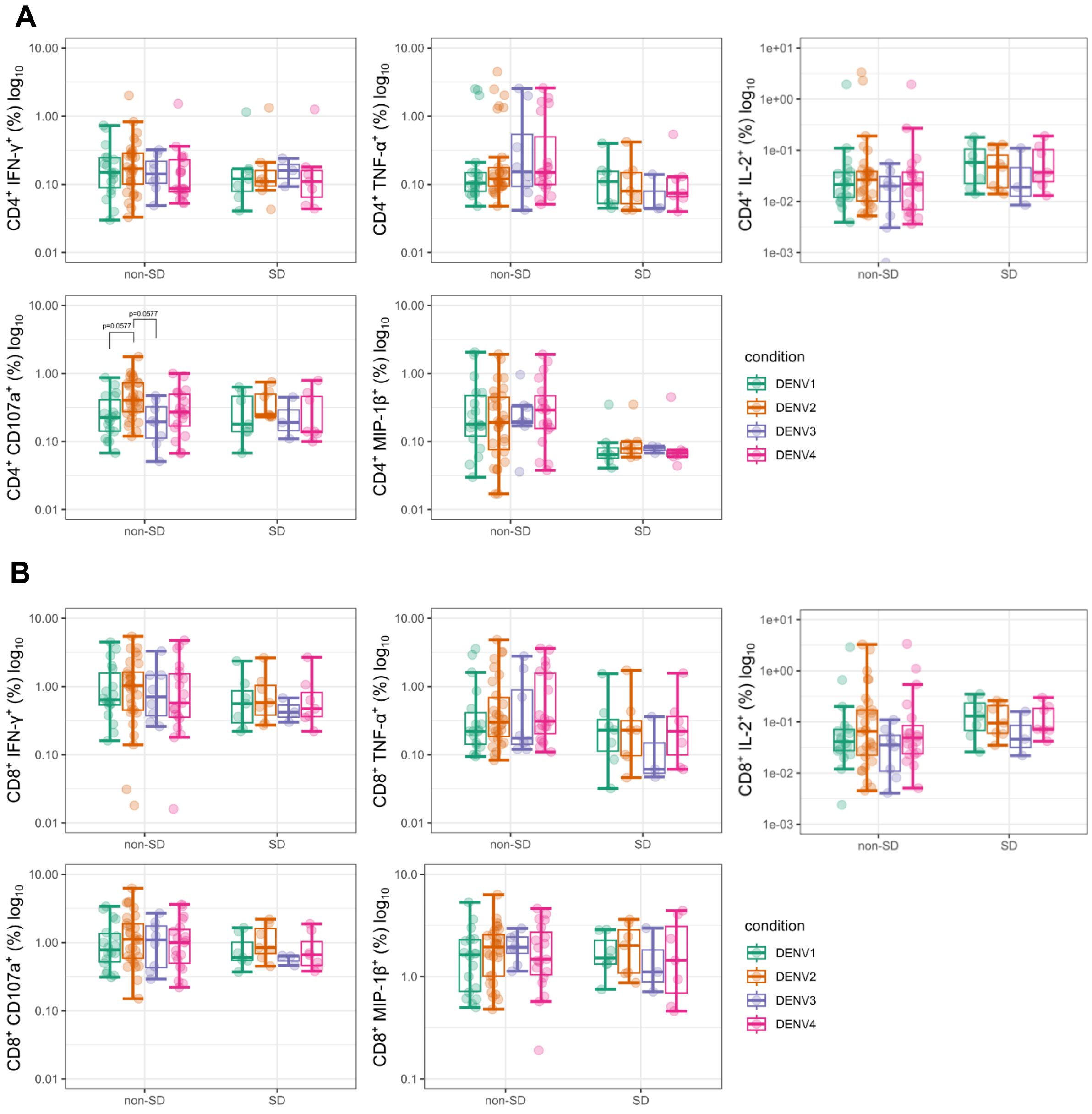


**Fig.S2. DENV NS3-specific T-cell response.** Single cytokine response by **(A)** CD4⁺ and **(B)** CD8⁺ T-cells in non-SD and SD patients at T2 (N=37) following NS3 DENV1-4 peptide stimulation.

### Supplementary Figure S3

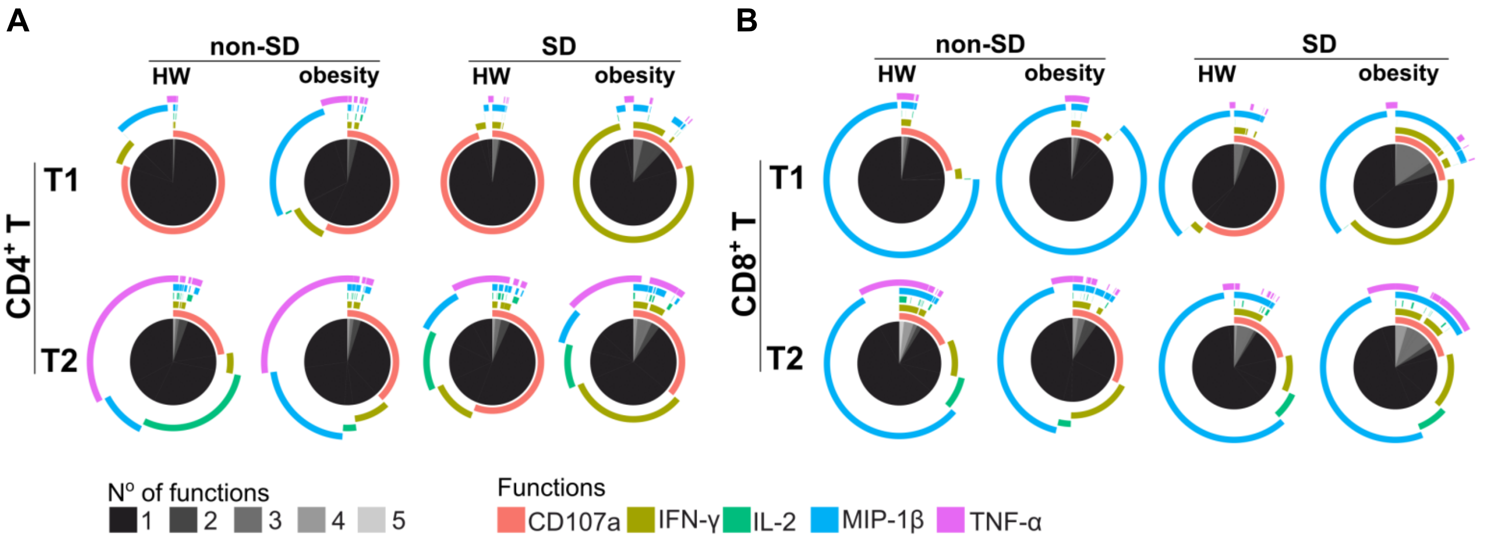


**Fig.S3. Assessment of DENV2-specific T-cell function during DENV2 infection.** Pie charts showing the number of functions simultaneously exhibited by **(A)** $CD4^+$ and **(B)** $CD8^+$ T-cells following NS3 DENV2 peptide stimulation. The different shades of grey represent the range of 1-5 functions, the outer arcs indicate the specific functions (IFN-γ/TNF-α/IL-2/MIP-1β/CD107a) defined by Boolean gating.

### Supplementary Figure S4

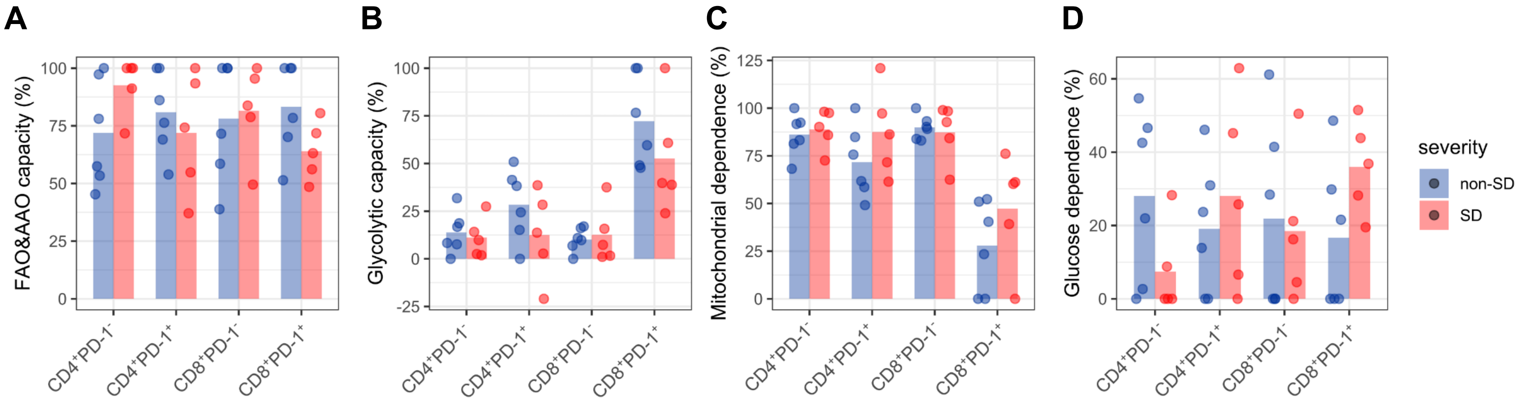


**Fig.S4. Metabolic activity measured by SCENITH. (A)** Fatty acid and amino acid oxidation (FAO&AAO) capacity, **(B)** glycolytic capacity, **(C)** mitochondrial dependence, and **(D)** glucose dependence. SCENITH was performed and capacities and dependencies were calculated according to Arguello et al.[32] and Luscombe et al.[33].

Supplementary Figure S5

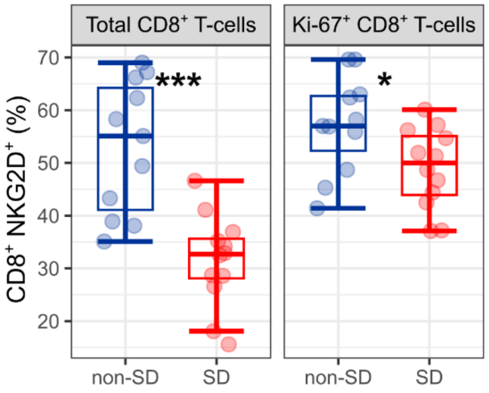


**Fig.S5. Impaired phenotype of $CD8^+$ T-cells in SD.** Frequency of total or Ki-67$^+$ $CD8^+$ T-cells expressing NKG2D in non-SD and SD patient group.

### Supplementary Figure S6

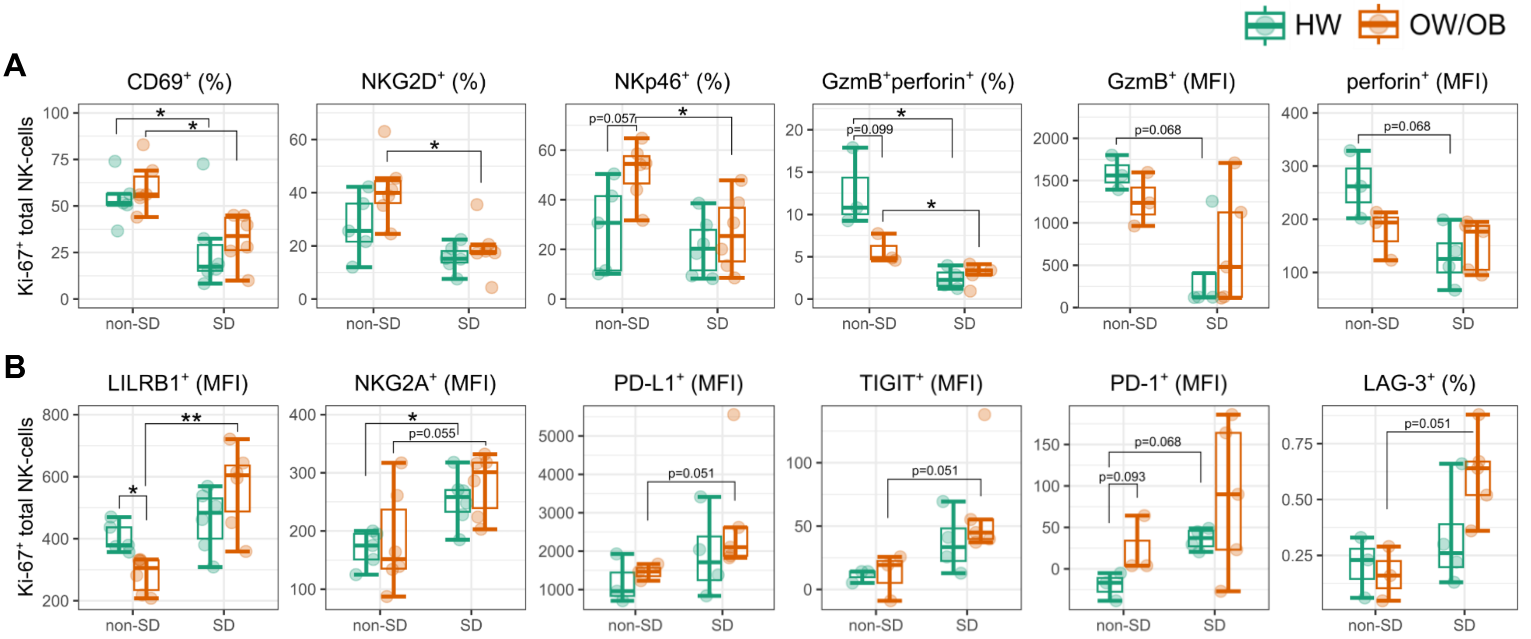


**Figure S6. Impaired phenotype of Ki-67⁺ total NK cells in SD.** Expression of **(A)** activating and **(B)** inhibitory receptors by Ki-67⁺ total NK-cells in HW (green) and OW/OB (orange) non-SD and SD patients.

Supplementary Figure S7

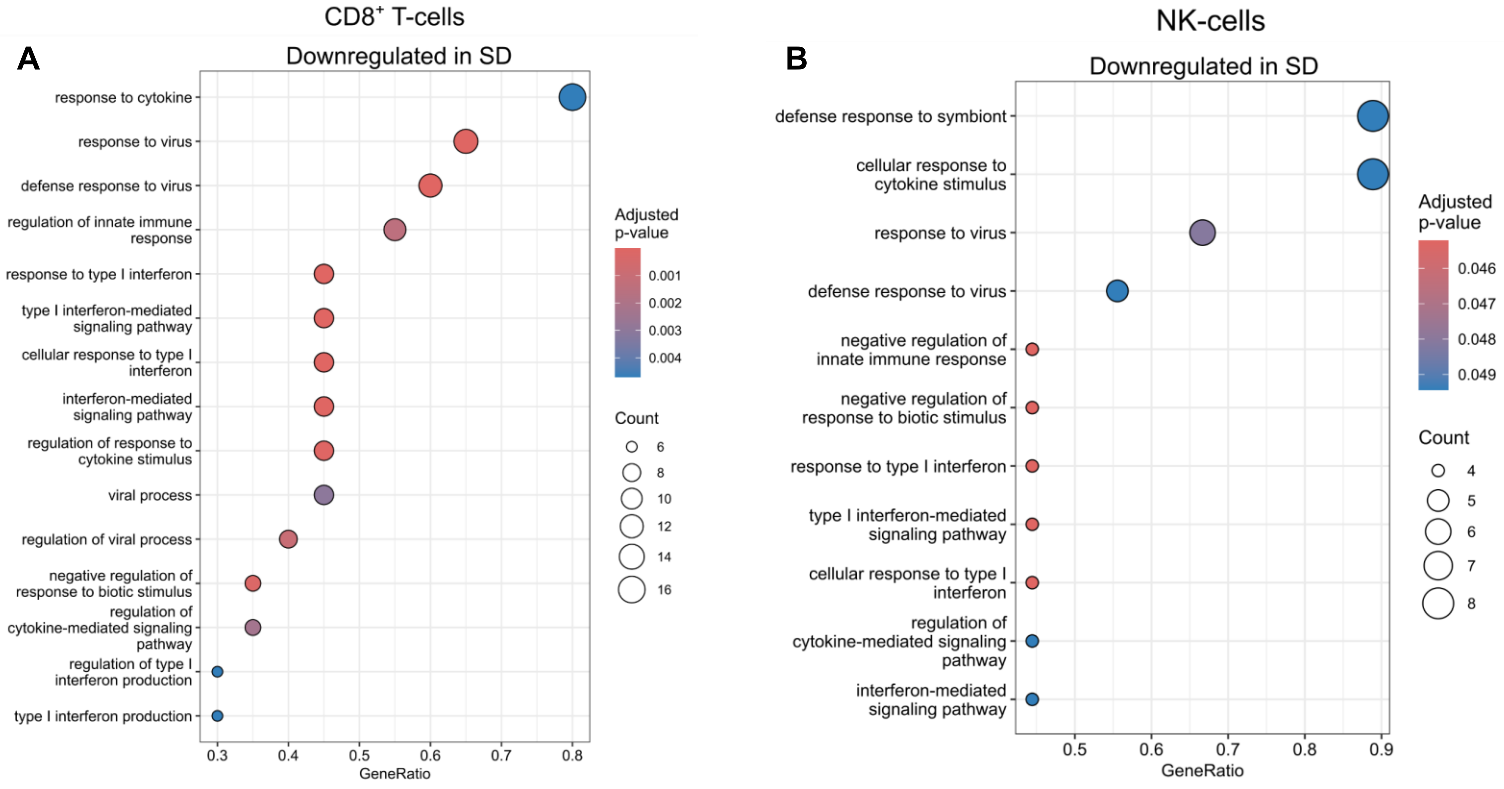


**Fig.S7. Impaired type I-IFN responses in SD.** Over-representation analyses of genes significantly downregulated in SD vs non-SD in CD8$^+$ T-cells **(A)** and NK-cells **(B)**. All significant or top 15 non-redundant gene ontology terms and associated BH adjusted p-values are shown. Count=number of differentially expressed genes (DEGs) in the gene set/pathway. GeneRatio=fraction of DEGs in the gene set. Data from N=24 dengue patients (N=12 non-SD; N=12 SD).

### Supplementary Figure S8

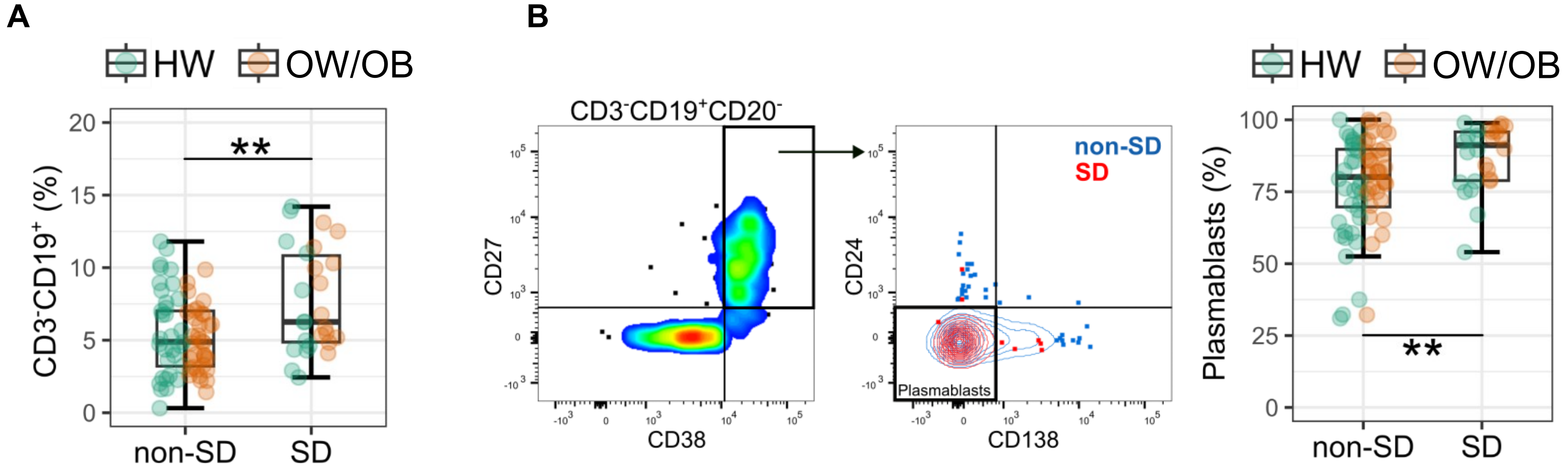


**Fig.S8. Increased frequency of B-cells and plasmablasts in SD. (A)** Frequency of $CD3^-CD19^+$ **(B)** Gating strategy and frequency of plasmablasts in non-SD and SD patients [HW (green) and OW/OB (orange)]. Data from N=94 dengue patients at T2 (N=68 non-SD; N=26 SD).

### Supplementary Table S1

**Table S1.** Summary table of clinical information at the admission time point.

| Severity | | Non-SD (n=94) | SD (n=30) |
|---|---|---|---|
| Age (median) | | 16 | 15 |
| Age group | Children (10-18y) | 56 | 26 |
| | Adult (19-30y) | 38 | 4 |
| Sex | Female | 23 | 7 |
| | Male | 71 | 23 |
| BMI group | Healthy weight | 44 | 16 |
| | Overweight/obesity | 50 | 14 |
| Serotype | DENV1 | 18 | 4 |
| | DENV2 | 65 | 21 |
| | DENV4 | 5 | 2 |
| | NEG | 6 | 3 |
| Infection | Primary | 5 | 0 |
| | Secondary | 89 | 29 |
| | N/A | 0 | 1 |
| Day of illness (enrolment) | Day 1 | 1 | 0 |
| | Day 2 | 22 | 2 |
| | Day 3 | 58 | 6 |
| | Day 4 | 13 | 5 |
| | Day 5 | 0 | 15 |
| | Day 6 | 0 | 1 |
| | Median (IQR) | 3 (3-3) | 5 (3-5) |
| Plasma leakage grade | Grade 0 | 59 | 1 |
| | Grade 1 | 35 | 0 |
| | Grade 2 | 0 | 29 |
| Warning signs | w/o | 14 | 0 |
| | w/ | 80 | 0 |
| | severe | 0 | 30 |

### Supplementary Table S2

**Table S2.** List of downregulated genes (all cell types combined) in SD patient group.

| gene | avg_log2FC | pct.1 | pct.2 | p_val_adj |
|---|---|---|---|---|
| *IFI27* | -0.00124 | 0.22 | 0.148 | 3.265976844258429e-5 |
| *TMEM97* | -0.02395 | 0.043 | 0.021 | 0.047492852504465576 |
| *IFNAR2* | -0.05297 | 0.219 | 0.158 | 0.008779 |
| *JUND* | -0.06451 | 0.955 | 0.963 | 0.007606 |
| *IRF4* | -0.11162 | 0.175 | 0.087 | 2.4999570364731466e-11 |
| *NFKBIA* | -0.12364 | 0.884 | 0.861 | 0.043724926494393586 |
| *CASP3* | -0.15132 | 0.186 | 0.112 | 1.1293082780582184e-6 |
| *PTPRC* | -0.15418 | 0.868 | 0.883 | 0.011105678841967843 |
| *CD37* | -0.19401 | 0.856 | 0.849 | 7.484156367918458e-5 |
| *CD3E* | -0.21027 | 0.564 | 0.629 | 0.005373 |
| *S100A11* | -0.2116 | 0.523 | 0.552 | 0.029512116651518212 |
| *CD69* | -0.22732 | 0.702 | 0.793 | 2.1503404924470268e-10 |
| *HLA-A* | -0.23046 | 1 | 1 | 5.301888946764544e-25 |
| *IFNAR1* | -0.24019 | 0.22 | 0.152 | 9.497397999920781e-4 |
| *JUNB* | -0.24219 | 0.744 | 0.788 | 7.291662743978101e-10 |
| *FCER1G* | -0.2482 | 0.245 | 0.303 | 0.025279229697721225 |
| *TRAC* | -0.25651 | 0.578 | 0.621 | 0.015588511421063099 |
| *ZFP36* | -0.25987 | 0.936 | 0.918 | 1.1077185424434842e-7 |
| *DUSP1* | -0.27983 | 0.947 | 0.965 | 3.645380249540277e-15 |
| *IL12A* | -0.29093 | 0.043 | 0.016 | 3.6926251530861894e-4 |
| *ID2* | -0.3081 | 0.48 | 0.547 | 1.35337097025562e-5 |
| *ITK* | -0.30858 | 0.42 | 0.461 | 0.024414898712325012 |
| *FYB* | -0.31729 | 0.57 | 0.66 | 2.4474485300143227e-8 |
| *SELL* | -0.32676 | 0.704 | 0.679 | 0.017384258538871034 |
| *SELPLG* | -0.3313 | 0.648 | 0.652 | 7.577726520062362e-4 |
| *IL2RG* | -0.33721 | 0.748 | 0.77 | 5.213397780188954e-7 |
| *FOS* | -0.36184 | 0.94 | 0.956 | 4.512322080542571e-26 |
| *LAT* | -0.36835 | 0.407 | 0.453 | 0.009415 |
| *IFITM2* | -0.37403 | 0.583 | 0.625 | 5.483156484053746e-6 |
| *CD4* | -0.39184 | 0.307 | 0.382 | 3.798707016645441e-5 |
| *LEF1* | -0.40127 | 0.321 | 0.368 | 0.026550630042775445 |
| *NR4A1* | -0.41058 | 0.239 | 0.289 | 0.017573494067437204 |
| *CD48* | -0.42706 | 0.861 | 0.877 | 8.587165353205064e-23 |
| *FAM65B* | -0.45292 | 0.498 | 0.595 | 1.4142462286946276e-11 |
| *BCL2A1* | -0.48191 | 0.21 | 0.267 | 0.001042 |
| *GIMAP5* | -0.52212 | 0.212 | 0.293 | 7.852835086515958e-7 |
| *AQP9* | -0.5235 | 0.086 | 0.123 | 0.040864 |
| *IL7R* | -0.55511 | 0.367 | 0.458 | 7.025623472782824e-10 |
| *IFITM3* | -0.60652 | 0.492 | 0.564 | 1.0801164855794629e-11 |
| *CCL4* | -0.60747 | 0.312 | 0.374 | 7.234502808786648e-5 |
| *LAP3* | -0.66574 | 0.523 | 0.561 | 3.5343273866115514e-13 |
| *TRAT1* | -0.68496 | 0.171 | 0.219 | 0.003936 |
| *LGALS9* | -0.72835 | 0.328 | 0.447 | 2.1860033174681364e-18 |
| *IFITM1* | -0.73664 | 0.229 | 0.315 | 9.9328750892645e-10 |
| *ADAR* | -0.75944 | 0.731 | 0.798 | 6.700888579606053e-42 |
| *IL1B* | -0.76319 | 0.158 | 0.198 | 0.028730273666547828 |
| *IRF9* | -0.77291 | 0.424 | 0.485 | 1.409518891603742e-11 |
| *GIMAP2* | -0.78155 | 0.384 | 0.491 | 8.068722309019342e-16 |
| *TNF* | -0.7954 | 0.209 | 0.267 | 4.5750764698257245e-5 |
| *EGR3* | -0.81286 | 0.061 | 0.11 | 4.184586522246943e-6 |
| *IFI16* | -0.82645 | 0.467 | 0.572 | 2.3877345424736146e-26 |
| *ZBP1* | -0.83209 | 0.318 | 0.397 | 1.871962521619138e-11 |
| *OSM* | -0.86268 | 0.053 | 0.091 | 3.444712150641305e-4 |
| *CX3CR1* | -0.8978 | 0.225 | 0.279 | 1.6653041519554325e-4 |
| *GBP5* | -0.90265 | 0.21 | 0.284 | 1.8062972377737617e-8 |
| *SOCS2* | -0.90943 | 0.056 | 0.091 | 0.003775 |
| *STAT2* | -1.01016 | 0.448 | 0.558 | 4.054749159525832e-32 |
| *MT2A* | -1.11326 | 0.502 | 0.713 | 1.567286568594581e-77 |
| *OAS1* | -1.13365 | 0.183 | 0.256 | 7.883789892343615e-10 |
| *IL1RN* | -1.17056 | 0.089 | 0.143 | 1.3034845260051035e-6 |
| *IRF7* | -1.17253 | 0.296 | 0.423 | 7.242802520225324e-28 |
| *IFIH1* | -1.22886 | 0.16 | 0.277 | 2.120019094844042e-22 |
| *IER3* | -1.37956 | 0.131 | 0.215 | 1.918248955580432e-13 |
| *USP18* | -1.3997 | 0.222 | 0.363 | 2.2766887842634936e-31 |
| *STAT1* | -1.44667 | 0.157 | 0.237 | 6.545196682277028e-12 |
| *DDX58* | -1.49001 | 0.04 | 0.091 | 2.667602827404134e-9 |
| *TNFSF10* | -1.55967 | 0.361 | 0.459 | 1.4634121293889796e-30 |
| *ISG15* | -1.57966 | 0.419 | 0.655 | 1.3604247083280757e-96 |
| *LAMP3* | -1.64389 | 0.056 | 0.151 | 1.1279257507540959e-23 |
| *CXCL11* | -1.94217 | 0.007 | 0.023 | 0.005333 |
| *CCL2* | -2.01643 | 0.028 | 0.108 | 2.0329045335334454e-25 |
| *IL1A* | -2.33252 | 0.011 | 0.028 | 0.026750095822707296 |
| *CXCL10* | -2.7476 | 0.027 | 0.125 | 9.250255353097799e-36 |

### Supplementary Table S3

**Table S3.** List of custom genes from BD Rhapsody gene panel.

| Gene | | |
|---|---|---|
| *ACE2* | *IKZF3* | *OAS1* |
| *ADAR* | *IL10* | *OSM* |
| *ADGRG1* | *IL17A* | *PDGFA* |
| *AHNAK* | *IL17RE* | *PDGFB* |
| *AHR* | *IL18BP* | *PGF* |
| *ANXA1* | *IL1A* | *PTPN22* |
| *AQP3* | *IL1R1* | *REL* |
| *AREG* | *IL21R* | *RUNX1* |
| *BACH1* | *IL23A* | *RUNX2* |
| *BATF* | *IL26* | *S100A11* |
| *BATF3* | *IL2RG* | *S100A8* |
| *BCL2L1* | *IL6R* | *S1PR1* |
| *BTG2* | *IL6ST* | *SATB1* |
| *CASP1* | *IL9R* | *SGK1* |
| *CASP3* | *IRF1* | *SIT1* |
| *CASP8* | *IRF3* | *SLAMF1* |
| *CCR6* | *IRF7* | *SLC16A1* |
| *CD40LG* | *IRF9* | *SLC4A10* |
| *CD68* | *ISG15* | *SMAD3* |
| *CEBPD* | *ITGA1* | *SOCS1* |
| *CREM* | *ITGAL* | *SOCS2* |
| *CSF1* | *ITK* | *SOCS3* |
| *DDIT3* | *JUND* | *STAT2* |
| *DDX58* | *KIR2DL4* | *STING1* |
| *DLL4* | *KLRC2* | *TGFA* |
| *FGF1* | *KLRD1* | *THEMIS* |
| *FGF22* | *LAYN* | *TNFAIP3* |
| *FGF9* | *LILRB1* | *TNFRSF18* |
| *FOS* | *LILRB2* | *TNFRSF1A* |
| *FOSL2* | *MAF* | *TNFRSF1B* |
| *FOXO3* | *MAPK1* | *TNFSF9* |
| *FURIN* | *MMP25* | *TOX* |
| *GATA3* | *MMP28* | *TRBV28* |
| *GBP5* | *MR1* | *TRDV2* |
| *GPR171* | *MT2A* | *USP18* |
| *GPR183* | *MYB* | *VCAM1* |
| *GPR65* | *NFAT5* | *VDR* |
| *GZMM* | *NFATC1* | *WNT1* |
| *HIF1A* | *NFATC2* | *WNT10A* |
| *HOPX* | *NFKB1* | *WNT3A* |
| *ID2* | *NFKB2* | *WNT6* |
| *ID3* | *NFKBIA* | *XCL1* |
| *IFI16* | *NFKBIZ* | *XCL2* |
| *IFI27* | *NOTCH1* | *ZBP1* |
| *IFIH1* | *NR1D1* | *ZBTB32* |
| *IFITM1* | *NR3C1* | *ZEB2* |
| *IFNAR1* | *NR4A1* | *ZFP36* |
| *IFNAR2* | *NR4A2* | |
| *IFNGR2* | *NR4A3* | |
